# Regulation of neural gene expression by estrogen receptor alpha

**DOI:** 10.1101/2020.10.21.349290

**Authors:** Bruno Gegenhuber, Melody V. Wu, Robert Bronstein, Jessica Tollkuhn

**Affiliations:** Cold Spring Harbor Laboratory, Cold Spring Harbor, NY 11724, USA; Cold Spring Harbor Laboratory School of Biological Sciences, Cold Spring Harbor, NY 11724 USA

## Abstract

The transcription factor estrogen receptor α (ERα) is a principal regulator of sex differences in the vertebrate brain and can modulate mood, behavior, and energy balance in females and males. However, the genes regulated by ERα in the brain remain largely unknown. Here we reveal the genomic binding of ERα within a sexually dimorphic neural circuit that regulates social behaviors. We profiled gene expression and chromatin accessibility and show ERα induces a neurodevelopmental gene program in adulthood. We further demonstrate that ERα binds with Nuclear factor I X-type (Nfix) to regulate a male-biased gene expression program that initiates in early life. Our results reveal a neural strategy for ERα-mediated gene regulation and provide molecular targets that underlie estrogen’s effects on brain development, behavior, and disease.

## Introduction

Estrogen is the master regulator of sexual differentiation of the rodent brain. It acts on discrete populations of neurons to define sex differences in neural circuits that mediate innate behaviors (MacLusky and Naftolin, 1981; McEwen, 1981). During a perinatal critical period, circulating testosterone in males is converted to 17β-estradiol, the principal endogenous estrogen, in the brain by aromatase (MacLusky and Naftolin, 1981). Neural estradiol then acts through ERα to specify sex differences in cell survival and circuit connectivity, which enable distinct social behaviors in adult females and males (Li and Dulac, 2018; McCarthy, 2008). How transient exposure to estrogen organizes lasting sex differences remains a central question in neurobiology. Estrogen also provides neuroprotection against inflammation and neurodegeneration (Arevalo et al., 2015), enhances learning and memory (Galea et al., 2017), and improves symptoms of depression and schizophrenia (Kulkarni et al., 2019; Wharton et al., 2012). Over the last 40 years, mechanisms of transcriptional control by ERα have been characterized in peripheral tissues and breast cancer, leading to diagnostic and therapeutic breakthroughs (Carroll, 2016; Evans, 1988). However, the genes regulated by ERα in the brain have not been identified.

## Results

### Identification of direct ERα binding sites in the brain

Sex-typical behaviors are regulated by sexually dimorphic limbic pathways that encompass the primary sites of ERα expression in the brain (Li and Dulac, 2018; Simerly, 2002). We previously demonstrated that loss of ERα in inhibitory GABAergic neurons is sufficient to dysmasculinize the brain and behavior (Wu and Tollkuhn, 2017), and sought to determine the genomic targets of this receptor to understand how it specifies sex differences. To date, no studies have identified direct binding sites of a gonadal hormone receptor in the brain, as the low and sparse expression of these receptors prohibits the use of ChIP-seq, which typically requires millions of cells. To bypass this issue, we used CUT&RUN to profile ERα in the brain. We first optimized low-input ERα CUT&RUN in MCF7 breast cancer cells, and validated prior findings (Fig. S1). We then performed CUT&RUN in pooled tissue from sexually dimorphic ERα+ brain regions that are primarily GABAergic: the posterior bed nucleus of the stria terminalis (BNSTp), medial pre-optic area (MPOA), and posterior medial amygdala (MeAp) from either adult males or females (Fig. 1A) (Li and Dulac, 2018; Simerly, 2002; Wu and Tollkuhn, 2017). We treated gonadectomized females and males with estradiol benzoate (EB) or corn oil vehicle (Veh) for 4 hours to synchronize transient nuclear receptor binding and capture of target sites. With this approach, we detected 1930 EB-induced ERα peaks (Fig. 1B, S2, Data S1). The most enriched TF binding motifs in these peaks are ERβ (ESR2) and ERα (ESR1) (Fig. S2), both of which can be bound by ERα (Grober et al., 2011). An ERα footprint was present at the ESR1 motif in these peaks (Fig. 1C), demonstrating the specificity of the method. We also detected 185 ERα peaks specific to vehicle (Fig. S2), for which the top enriched motifs were CTCF and RFX5, suggesting ERα coordinates distinct chromatin interactions in the absence of ligand. Strikingly, we detect few sex differences in ERα binding (Fig. S2), indicating that adult female and male brains possess the same capacity to respond to estrogen signaling.

**Fig. 1.**
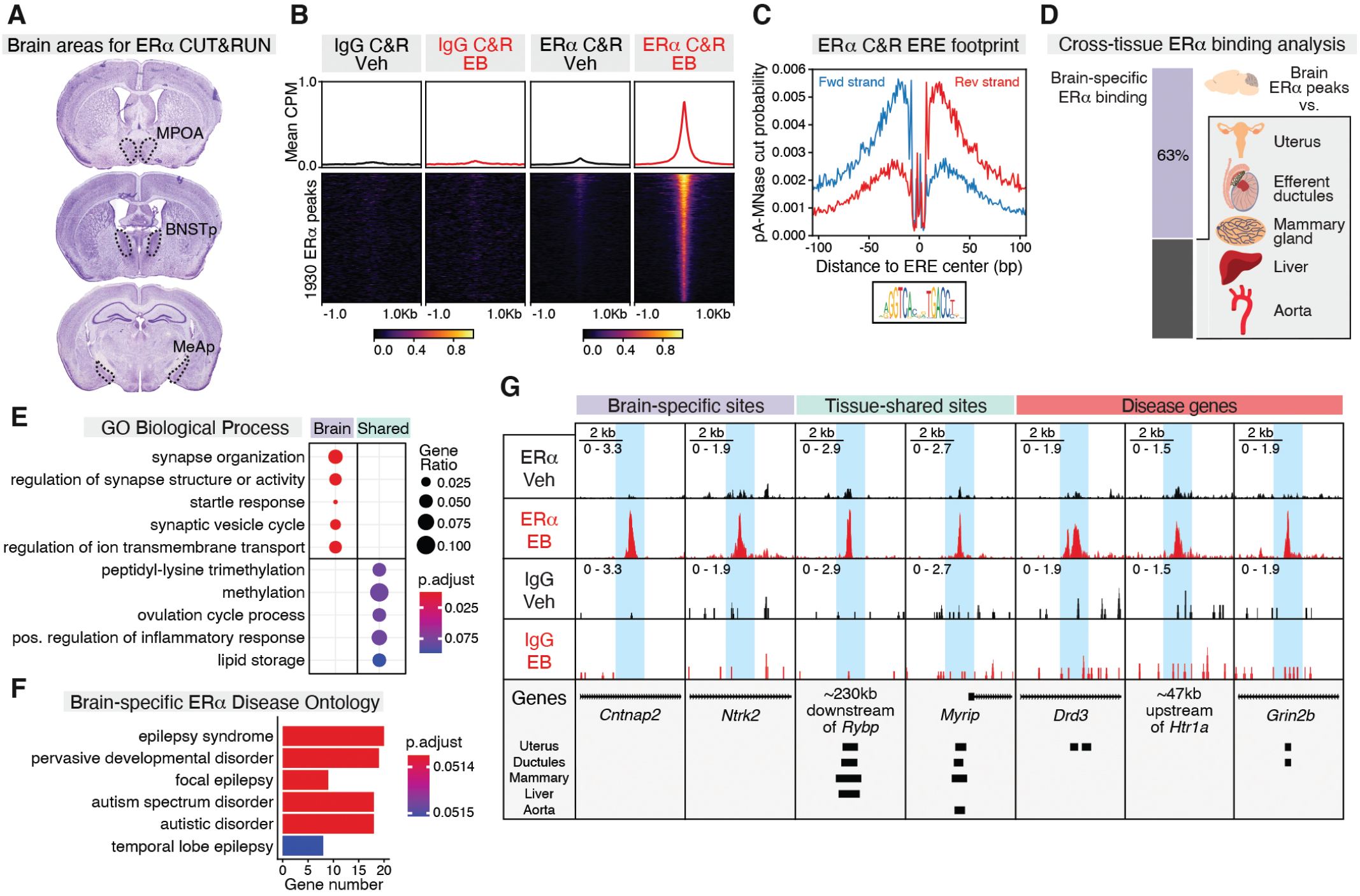
Genomic targets of ERα in sexually dimorphic neuronal populations. **(A)** Coronal sections of sexually dimorphic brain areas used for ERα CUT&RUN. **(B)** Heatmap of mean IgG and ERα CUT&RUN CPM +/−1Kb around EB-induced ERα CUT&RUN peaks (DiffBind edgeR, padj<0.1). Heatmap sorted by EB ERα CUT&RUN signal. **(C)** ESR1 motif footprint in ERα peaks (CUT&RUNTools). **(D)** Proportion of ERα peaks detected specifically in brain. **(E)** Top GO Biological Process terms associated with genes nearest to brain-specific or shared (≥4 other tissues) ERα peaks (clusterProfiler, padj<0.1). **(F)** Top Disease Ontology (DO) terms associated with genes nearest to brain-specific ERα peaks (DOSE, padj<0.1). **(G)** Example brain-specific (*Cntnap2*, *Ntrk2*), shared (*Rybp*, *Myrip*), and disease-associated (*Drd3*, *Htr1a*, *Grin2b*) ERα peaks.

We compared our ERα peaks to published ERα ChIP-seq data collected from peripheral mouse tissues, revealing that the majority of our ERα peaks are brain-specific (Fig. 1D). Genes associated with brain-specific peaks are enriched for synaptic and neurodevelopmental disease GO terms, including neurotransmitter receptors, neurotrophin receptors, and extracellular matrix genes (Fig. 1E, 1F, S3, Data S1). These direct ERα target genes account for many of the known phenotypic effects of estrogen. Examples include the serotonin receptor *Htr1a* and dopamine receptor *Drd3*, which may facilitate estrogen control of observed sex differences in depression, schizophrenia, and Parkinson’s disease symptoms (Fig. 1G) (Abel et al., 2010; Altemus et al., 2014; Meoni et al., 2020). The effects of estrogen on neuroprotection and synaptic plasticity are similar to those of the neurotrophin BDNF, and we find additional evidence supporting crosstalk between these two signaling systems (Carbone and Handa, 2013). ERα directly binds the gene for the BDNF receptor TrkB (*Ntrk2*) (Fig. 1G), as well as *Ntrk3*, an additional neurotrophin receptor. Autism shows a pronounced male bias, and male-specific early life exposure to estrogen could impart additional vulnerability to the effects of disruptive mutations in associated genes (Manoli and Tollkuhn, 2018). Accordingly, ERα is found on several high-confidence autism candidate genes including *Cntnap2, Grin2b, Scn2a*, and *Dyrk1a* (Fig 1G, Data S1) (Hoffman et al., 2016; Satterstrom et al., 2020). We also note that ERα binds downstream of *Ar*, the gene for androgen receptor (Data S1). Additional disease candidates include *App*, the gene for amyloid precursor protein, and *Bace2*, which encodes one of the two enzymes that cleaves App to form amyloid beta (Data S1). Collectively, our CUT&RUN data demonstrate estrogen can exert widespread control of neuronal function via ERα recruitment to a distinct set of genomic targets in the brain.

### Estrogen induction of gene expression and chromatin accessibility

To identify the functional effects of ERα genomic binding, we assessed estrogen-induced gene expression specifically within ERα neurons. To obtain increased specificity, we focused on a single GABAergic population, the BNSTp, which shows evolutionarily conserved sexual dimorphism (Allen and Gorski, 1990; Hines et al., 1992; Simerly, 2002; Welch et al., 2019; Zhou et al., 1995). In mice, this dimorphism requires ERα (Tsukahara et al., 2011). We performed translating ribosome affinity purification (TRAP) in microdissected BNSTp from *Esr1*^Cre/+^; *Rpl22*^HA/+^ females and males using the same treatment paradigm as above, followed by RNA-seq. We identified 358 genes with differential expression between EB and Veh treatment groups, including known estrogen-induced genes, such as *Pgr* and *Nrip1* (Fig 2A, Data S2). Pairwise comparisons by sex found 297 differential genes (Fig. S4, Data S2); however only a few genes were induced by estrogen specifically in one sex (Data S2), demonstrating that, as with ERα binding, female and male neurons possess the capacity to mount the same transcriptional response to estrogen signaling. Although estrogen treatment caused both induction and repression of gene expression, ERα binding was primarily localized at estrogen-induced genes (Fig. 2B).

**Fig. 2.**
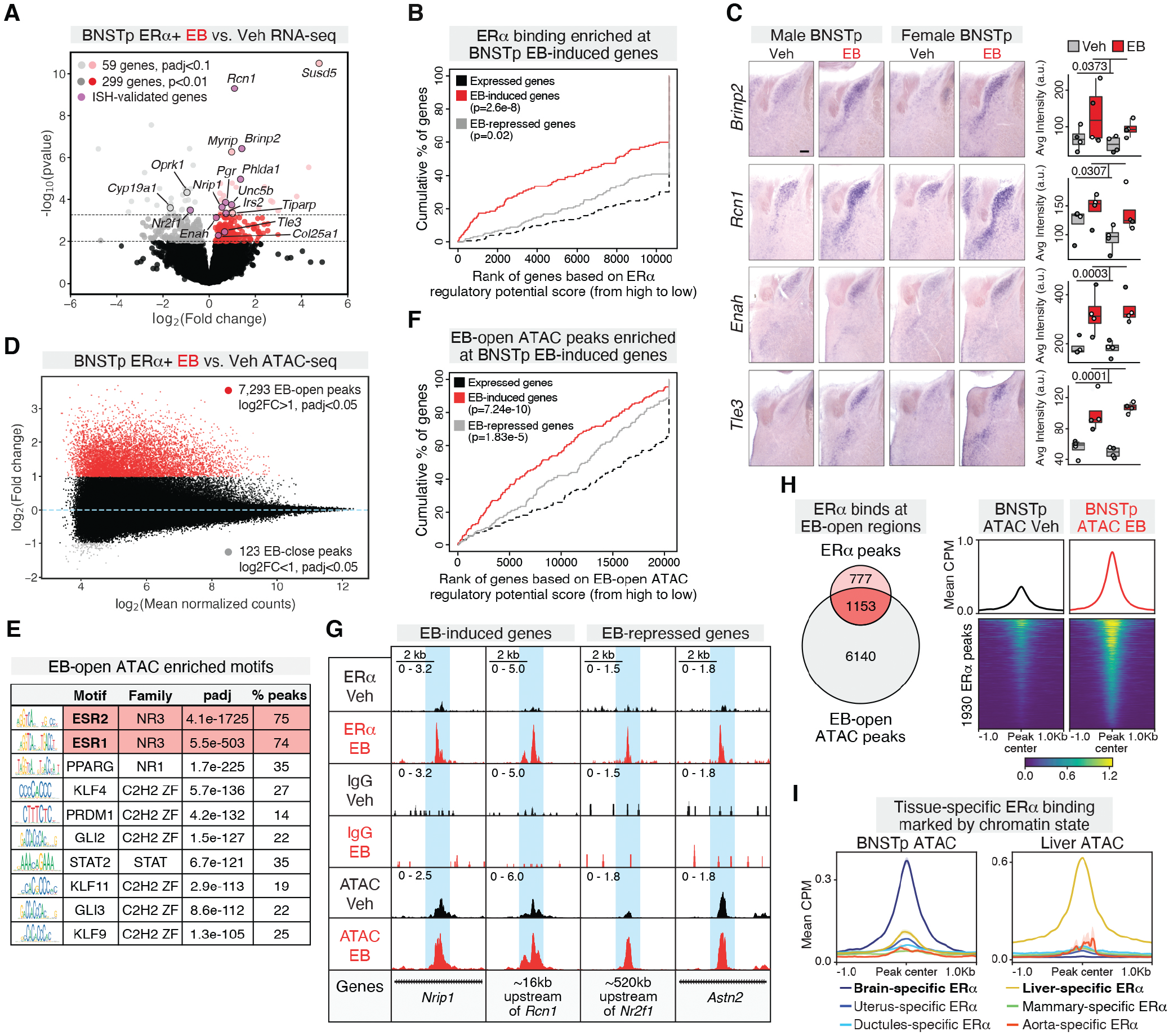
Estrogen regulation of gene expression and chromatin accessibility in BNSTp ERα+ cells. **(A)** Combined sex EB vs. Veh RNA-seq in BNSTp ERα+ cells; light grey and red dots (DESeq2, padj<0.1), dark grey and red dots (DESeq2, p<0.01), purple dots (ISH-validated). **(B)** BETA analysis of ERα peaks and differentially-expressed genes (DESeq2, p<0.01). p-values from K-S test. **(C)** ISH of select genes induced by EB in both sexes (2-way ANOVA: *Brinp2* p=0.0373, *Rcn1* p=0.0307, *Enah* p=0.0003, *Tle3* p=0.0001; n=4, scale=200um). **(D)** MA plot of EB-regulated ATAC-seq peaks in BNSTp ERα+ cells; red dots (DiffBind edgeR, log2FC>1, padj<0.05), grey dots (DiffBind edgeR, log2FC<-1, padj<0.05). **(E)** Top motifs enriched in EB-open ATAC-seq peaks (AME). % of peaks containing enriched motifs calculated with FIMO. **(F)** BETA analysis of EB-open ATAC sites and differentially-expressed genes (DESeq2, p<0.01). p-values from K-S test. **(G)** Example ERα peaks at estrogen-induced (*Nrip1*, *Rcn1*) and estrogen-repressed (*Nr2f1*, *Astn2*) genes. **(H)** (Left) Intersection of ERα peaks and EB-open ATAC peaks. (Right) Heatmap of mean ATAC CPM +/−1Kb around EB-induced ERα peaks. Heatmap sorted by EB ATAC CPM. **(I)** Mean+SD BNSTp vehicle ATAC CPM (left) and liver ATAC CPM (right) at tissue-specific, non-promoter ERα peaks.

We validated several of the top EB-induced genes by *in situ* hybridization (ISH) (Fig. 2C, Fig. S5), demonstrating genes involved in neuron wiring (*Brinp2, Unc5b, Enah*) and synaptic plasticity (*Rcn1, Irs2*) are robustly induced by estrogen in the adult BNSTp. ERα co-repressors *Tle3* and *Nrip1* increase expression with EB, while the co-activator *Nr2f1* is decreased, suggesting a negative transcriptional feedback mechanism (Fig. 2C, S5, Data S2). *Nr3c1*, the gene for glucocorticoid receptor showed increased expression in the female BNSTp and medial amygdala (Fig. S5), which may contribute to sex differences in stress responses (Bangasser and Wicks, 2017). EB-regulated genes showed variable expression patterns in other ERα+ brain regions, indicating region specificity in estrogen-regulated target genes (Fig. S5). Our results show that estrogen induces genes associated with axonogenesis and synaptic organization in the adult brain, suggesting potential candidates that contribute to sex-specific neuronal connectivity following perinatal estradiol signaling (Gu et al., 2003).

To determine the effects of ERα binding on chromatin state and detect additional estrogen-responsive loci, in particular those with transient binding, we performed ATAC-seq in FACS-purified BNSTp ERα+ neurons collected from *Esr1*^Cre/+^; *ROSA26*^CAG-Sun1-sfGFP-Myc/+^ mice using the same treatment paradigm (Fig. S6). Across sexes, we observed 7,293 chromatin regions that increase accessibility (EB-open) and 123 regions that decrease accessibility (EB-close) with EB treatment, demonstrating estrogen robustly induces chromatin opening in neurons (Fig. 2D, S6, Data S3). As with our RNA-seq, there are few sex differences in EB-responsive peaks (Fig. S6).

We find the effects of estrogen on chromatin state to be primarily mediated by direct actions of hormone receptors and not via other estrogen-mediated signaling pathways (Micevych and Kelly, 2012; Srivastava et al., 2013); 89% of EB-open ATAC peaks (6479/7293) contain either an ESR1 or ESR2 motif (Fig. 2E, S6). In contrast to our ERα binding data, EB-open ATAC peaks are localized at both EB-upregulated and -downregulated genes (Fig. 2F), demonstrating estrogen-induced chromatin opening can lead to both gene activation and repression. Example upregulated genes include ISH-validated targets such as *Rcn1* and *Nrip1*; downregulated genes include *Astn2*, a regulator of synaptic trafficking, and *Nr2f1* (Fig 2G). EB-open ATAC peaks associated with downregulated genes were enriched for ESR motifs (Fig. S6), suggesting additional ERα binding events undetected by CUT&RUN contribute to gene repression (Guertin et al., 2014). ~60% of estrogen-induced ERα peaks (1153/1930) overlap a robust, EB-open ATAC peak (Fig. 2H, S6), while the majority of remaining ERα peaks (521/777) overlapped genomic regions that exhibited a smaller change in accessibility (Fig. S6). Brain-specific ERα peaks are accessible in BNSTp, while ERα peaks specific to other tissues are not (Fig. 2I). Interestingly, brain-specific BNSTp-accessible ERα peaks contain weaker estrogen response elements (EREs) than tissue-shared ERα peaks, suggesting tissue-specific ERα binding patterns are facilitated by cooperativity with additional cofactors (Fig. S3). Importantly, our findings indicate differential exposure to estrogen in early life, or following puberty, does not constrain the chromatin landscape or potential for acute ERα regulation in adults.

### Nfix coordinates dynamic regulation of gene expression by ERα

ERα cooperates with different TFs across tissues to regulate gene expression, although no ERα interaction partners have yet been identified in the brain (Droog et al., 2016). Among the top enriched motifs in ERα peaks is the NFI monomer motif (Fig. 3A); NFI factors are known to cooperate with nuclear hormone receptors to regulate transcription (Wiench et al., 2011). We noted that the NFI family member Nfix is expressed in a discrete subset of BNSTp neurons (Fig 3A), and shows male-biased expression in BNSTp ERα+ neurons (Fig. S4). To determine whether ERα and Nfix share direct target genes, we profiled Nfix genomic binding in the BNSTp with CUT&RUN after *in vitro* validation (Fig. S7). We detected 32,578 Nfix peaks conserved across sex and treatment (Fig. S7). A small proportion of Nfix peaks varied by sex (158 male-biased, 99 female-biased) and treatment (524 EB-induced, 106 EB-reduced) (Fig. S7). The top enriched motifs at Nfix peaks were NFIX and CTCF motifs (Fig. S7), consistent with NFI factor enrichment at chromatin boundaries (Pjanic et al., 2013).

**Fig. 3.**
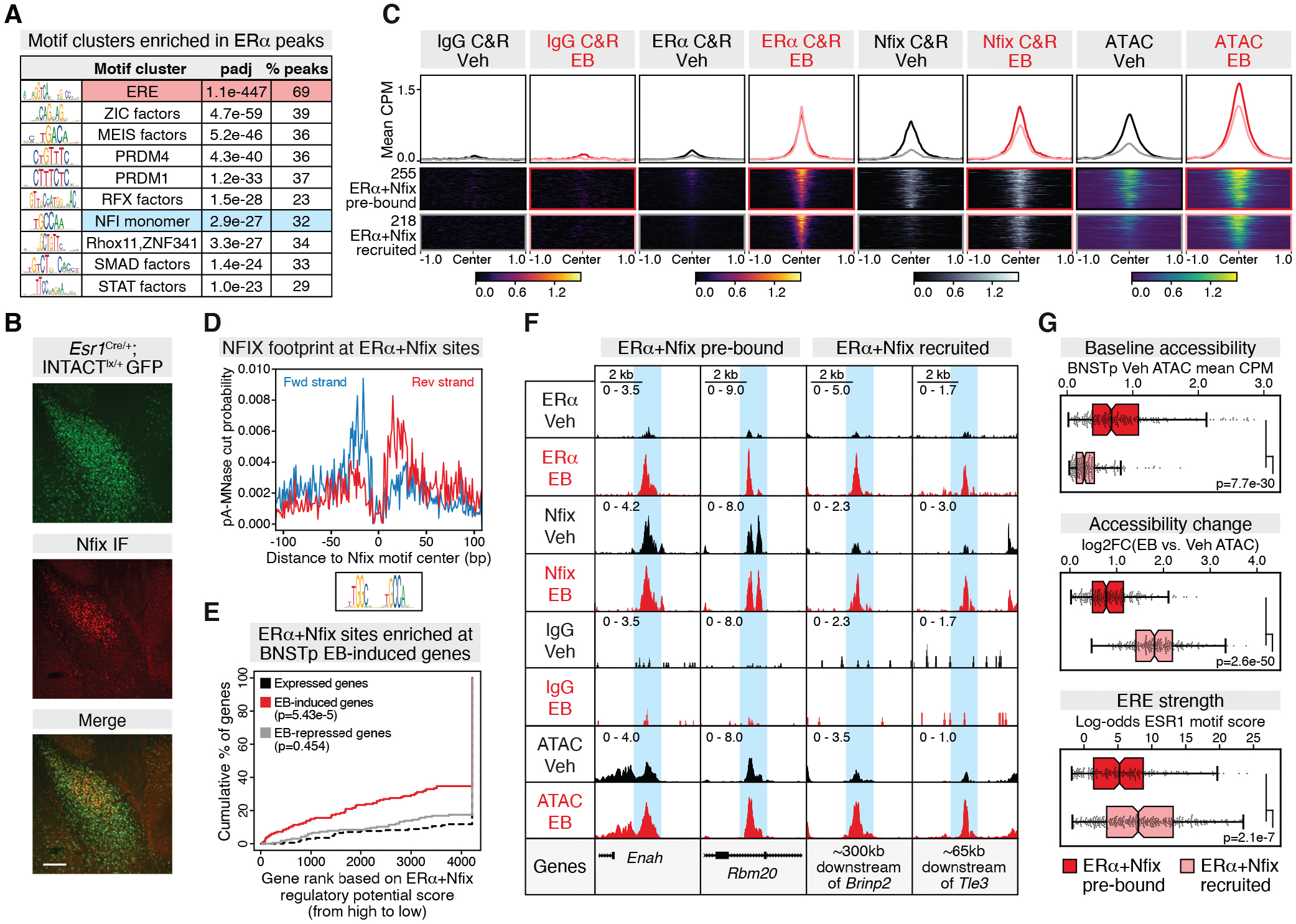
Nfix and ERα are bimodally recruited to estrogen-induced genes. **(A)** Top enriched JASPAR motif clusters (AME) in ERα peaks. % peaks containing enriched motif determined with FIMO. **(B)** Immunofluorescence staining for GFP and Nfix in an adult male *Esr1*^Cre/+^; *ROSA26*^CAG-Sun1-sfGFP-Myc/+^ animal (scale=100mm). **(C)** Mean IgG CUT&RUN, ERα CUT&RUN, Nfix CUT&RUN, and ATAC CPM +/−1Kb around 255 ERα+Nfix pre-bound peaks (dark lines) and 218 ERα+Nfix recruited peaks (light lines). Heatmap sorted by mean EB ERα CUT&RUN CPM. **(D)** NFIX dimer motif footprint at ERα+Nfix co-bound peaks (CUT&RUNTools). **(E)** BETA analysis of ERα+Nfix co-bound peaks and differential genes (DESeq2, p<0.01). p-value from K-S test. **(F)** Example ERα+Nfix pre-bound (*Enah*, *Rbm20*) and ERα+Nfix recruited (*Brinp2*, *Tle3*) peaks at estrogen-induced genes. **(G)** Comparison of (top) baseline BNSTp chromatin accessibility (Veh ATAC mean CPM), (middle) accessibility fold change, and (bottom) log-odds ESR1 motif score at ERα+Nfix pre-bound (dark red) and ERα+Nfix recruited peaks (light red). p-values from Mann-Whitney *U* test.

We investigated whether ERα binds with Nfix and detected a bimodal pattern of ERα and Nfix recruitment in response to estrogen (Archer et al., 1992). Although Nfix is present in only a subset of ERα neurons, 25% of EB-induced ERα binding sites overlap with Nfix occupancy (Fig. 3C). 255 ERα sites overlapped non-differential Nfix peaks, indicating that Nfix was pre-bound before ERα recruitment, and 218 additional ERα sites overlapped EB-induced Nfix peaks, demonstrating Nfix recruitment to chromatin in the presence of ligand (Fig. 3C, S7). We detected a footprint over the Nfix motif across both categories of ERα+Nfix co-bound sites, indicating physical Nfix occupancy (Fig. 3D). As with total ERα peaks, ERα+Nfix co-bound sites are enriched at EB-induced genes, including ISH-validated targets *Brinp2, Tle3*, and *Enah* (Fig. 3E-F). Finally, ERα+Nfix recruited sites were less accessible, had higher EB-induced accessibility, and had stronger EREs (Fig. 3G) than pre-bound sites, indicating ERα recruits Nfix to low-accessible regions through high-affinity motif binding and subsequent chromatin opening. Our results demonstrate ERα and Nfix regulate a distinct module of estrogen-responsive genes in the BNSTp.

### Sex differences in gene expression in the brain are associated with ERα genomic occupancy

As a master regulator, ERα organizes posterior BNST circuitry to enable male-typical behaviors in adulthood (Ogawa et al., 1997; Tsukahara et al., 2011; Wu and Tollkuhn, 2017). We hypothesized that ERα target genes have a persistent sex difference in expression following the perinatal estrogen surge, suggesting the timing of ERα function contributes to sexual differentiation of the brain. Prior studies have detected only minor sex differences in brain gene expression, and the factors driving these sex differences remain unclear. To identify sex differences in BNST gene expression, we re-analyzed a single-nucleus RNA-seq (snRNA-seq) dataset (Welch et al., 2019), consisting of 76,693 neurons across both sexes. We performed pseudo-bulk differential expression analysis within transcriptionally-defined BNST neuron types. Across 40 snRNA-seq clusters, nearly all sexually dimorphic genes were detected in *Esr1*-expressing clusters (Fig. 4A, S8), revealing ERα expression is predictive of sex differences in gene expression in the BNST.

**Fig. 4.**
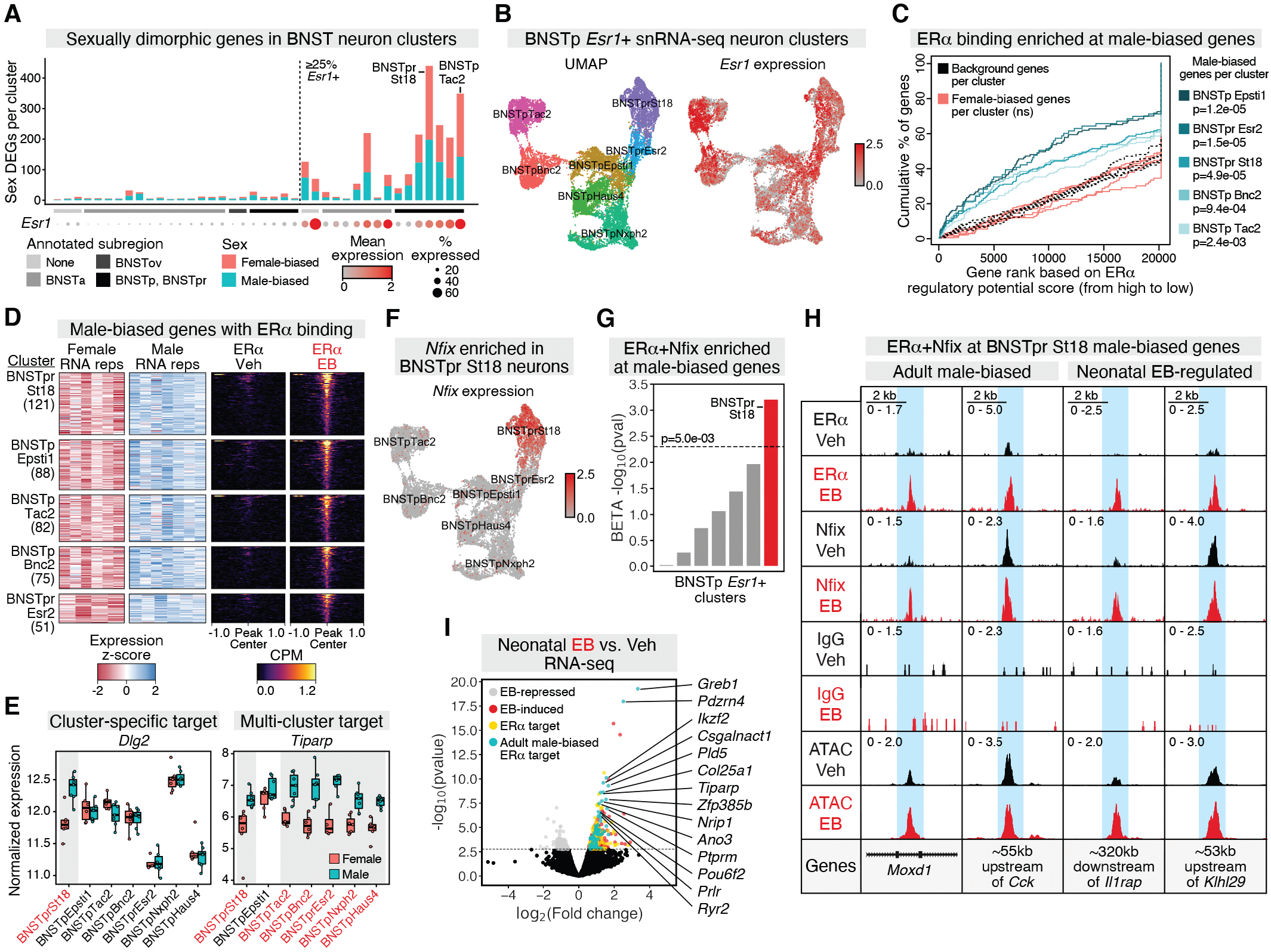
ERα binds extensively at male-biased genes in the BNST. **(A)** Number of sexually dimorphic genes (DEGs) per cluster (DESeq2, padj<0.1) across BNST snRNA-seq clusters. Clusters segregated by *Esr1* expression, grouped by BNST subregion, and sorted by % *Esr1*+ nuclei per cluster. **(B)** (Left) UMAP of *Esr1*+ BNSTp clusters. (Right) Normalized *Esr1* counts per nucleus. **(C)** BETA analysis of ERα peaks and sex DEGs detected in *Esr1*+ clusters. p-values from K-S test, ns=not significant. **(D)** (Left) Heatmap showing z-scaled, normalized expression of male-biased ERα target genes for the 5 clusters with enriched ERα binding. (Right) Mean ERα CUT&RUN CPM +/−1Kb around the nearest ERα peak to male-biased genes in each cluster. Heatmaps sorted by EB ERα CUT&RUN CPM. **(E)** Boxplots showing normalized expression of *Dlg2* and *Tiparp*. Red=clusters with differential expression. **(F)** Normalized *Nfix* counts per nucleus in *Esr1*+ clusters. **(G)** Barplot showing BETA analysis −log10(p-val) for ERα+Nfix co-bound peaks and male-biased genes in *Esr1*+ clusters. Dashed line=K-S test cutoff. **(H)** Example ERα+Nfix co-bound peaks at BNSTpr St18 male-biased genes and BNSTpr St18 male-biased genes regulated by neonatal EB. **(I)** Neonatal female EB vs. Veh RNA-seq in BNST ERα+ nuclei; grey, red dots (DESeq2, padj<0.05); ERα target genes not male-biased (gold dots) or male-biased (cyan dots) in adult BNSTp.

Among the 16 *Esr1*-expressing snRNA-seq clusters, we observed a higher number of sexually dimorphic genes in clusters annotated to the BNSTp, including the principal nucleus (BNSTpr), than the anterior BNST (Fig. 4A). We integrated our ERα binding data with sexually dimorphic genes detected in BNSTp clusters to determine whether they are direct ERα targets. Male-biased, but not female-biased, genes in 5 of the 7 BNSTp *Esr1*-expressing clusters were significantly enriched for ERα binding sites (Fig. 4C). Male-biased ERα target genes include 5 of 9 genes (*Cartpt*, *Cckar*, *Ecel1*, *Greb1*, *Sytl4*) previously validated in the medial BNSTp (Xu et al., 2012) and over 200 novel genes (Fig. 4D, Data S4, S5). The majority of these genes displayed male-biased expression in specific clusters (e.g., *Dlg2*), whereas a small proportion of genes were male-biased across multiple clusters (e.g., *Tiparp*) (Fig. 4E, S8). EB-open ATAC peaks were also enriched at genes that are male-biased in the BNSTp (Fig. S8). Together these results reveal estrogen-responsive loci as specifically enriched at BNSTp male-biased genes. As these loci were detected across both sexes, the results also indicate females retain the capacity for male-typical gene expression in adulthood. The mechanistic basis of female-biased gene expression remains unclear. These genes may be persistently downregulated in the neonatal male brain by transient ERα binding events (Guertin et al., 2014). Alternatively, ERα binding events that occur throughout the estrous cycle may not be captured following removal of hormones in adult females.

Given the observed role of Nfix in ERα genomic binding, we examined *Nfix* expression across BNSTp *Esr1*-expressing clusters. We detected specific enrichment of *Nfix* in BNSTpr St18 neurons (Fig. 4F, S9), consistent with our histologic data (Fig. 3B, Fig. S9). Across *Esr1*-expressing BNSTp clusters, we observed significant enrichment of ERα+Nfix co-binding only at male-biased genes in BNSTpr St18 neurons (Fig. 4G), with 55% of male-biased ERα target genes (66/121) in this cluster associating with an ERα+Nfix co-bound site (Fig. S9). Example co-bound targets include *Moxd1* - a marker of BNSTpr sexual dimorphism (Tsuneoka et al., 2017) and the neuropeptide *Cck* (Fig. 4H). Two of the top genes enriched in BNSTpr St18 neurons relative to other *Esr1*-expressing clusters, *Moxd1* and *Cplx3* (Fig. S9), were previously identified as markers of a scRNA-seq neuron cluster in the sexually dimorphic nucleus of the POA (SDN-POA) (i20:Gal/Moxd1) (Moffitt et al., 2018). Using integrated clustering, we observed that BNSTpr St18 neurons co-cluster with i20:Gal/Moxd1 neurons and that both populations are enriched for *Nfix* (Fig. S9), suggesting ERα and Nfix coordinate a male-biased transcriptional program across two sexually dimorphic brain regions.

To determine whether male-biased ERα target genes in the adult BNSTp are regulated by neonatal estrogen, we treated female mice at birth with EB or Veh and performed RNA-seq on FACS-purified BNST ERα+ nuclei four days later (Fig. 4I, Data S6). As in the adult brain, ERα binding was significantly enriched at neonatal EB-induced genes (208/359) but not EB-repressed genes (Fig. S10). We find that 33% of neonatal ERα target genes (69/208) are male-biased in the adult BNSTp (Fig. 4I, Data S5), consistent with the hypothesis that early life estrogen signaling organizes a lasting male-typical gene expression program (Gegenhuber and Tollkuhn, 2019; MacLusky and Naftolin, 1981). These genes likely specify sex differences in cell survival and connectivity to facilitate masculinized behavioral displays in adulthood, although the nature of the mechanism that canalizes male-biased expression remains an open question. Nevertheless, our findings show that ERα+ neurons in adults of both sexes have similar potential to respond to estrogen, although the functional effects of these genes are constrained by developmentally-specified circuit architecture. Such plasticity is recapitulated at the behavioral level: hormonal, circuit, and genetic manipulations consistently demonstrate that both sexes have the potential to display similar sex-typical behaviors (Edwards and Burge, 1971; Hashikawa et al., 2017; Kimchi et al., 2007; Wei et al., 2018). Collectively, these findings implicate hormonal regulation of gene expression in inhibitory neurons as the principal driver of the execution of sex differences in behavior.

## Discussion

Estrogen regulation of genes associated with neurological and psychiatric conditions may contribute to sex differences in the incidence and etiology of such diseases. Males are more often diagnosed with childhood-onset disorders including autism, ADHD, and dyslexia, while females have increased incidence of adolescent-onset mood and anxiety disorders. The onset of these disorders often follows sex-specific hormonal windows: men experience early-life exposure to testosterone or neural estradiol in the second trimester of gestation, while women undergo fluctuations in estrogens and progestins following puberty. Neurodegenerative diseases also show a sex-bias: men are more likely to develop Parkinson’s disease, and women are twice as likely to have Alzheimer’s. We expect many of the estrogen target genes and loci described here can contribute to susceptibility or resilience to developing disease, and that other genetic or environmental factors can intersect with mutations in these target loci to produce sex-specific outcomes.

Our identification of brain ERα targets and cofactors could facilitate the development of improved selective estrogen receptor modulators (SERMs) to provide the therapeutic benefits of estrogen on cognition, mood, and sleep, without increased risk of reproductive cancers and coronary artery disease. More broadly, we outline an approach to functionally connect a TF to innate behavior phenotypes via identification of its target genes in relevant neuronal populations. As single-cell methods continue to reveal TFs that define neuronal identity, the next step will be to determine how the direct targets of those factors contribute to the development and function of neural circuits.

## Acknowledgements

This work was performed with assistance from CSHL Shared Resources, including the Animal, Next Generation Genomics, Flow Cytometry, and Microscopy Core Facilities, which are supported by the Cancer Center Support Grant 5P30CA045508.

## Author contributions

BG carried out the ERα CUT&RUN and ATAC-seq experiments, and all bioinformatic analyses. MVW performed RNA-seq and ISH experiments. RB performed the Nfix CUT&RUN. BG and JT conceived the study, designed the experiments, and wrote the manuscript.

## Data and materials availability

All sequencing data deposited in GEO (GSE144718). All other data are available in the manuscript or the supplementary materials. Supplementary data files (Data S1-S6) will be provided upon request.

## Materials and Methods

### Animals

All mouse experiments were performed under strict guidelines set forth by the CSHL Institutional Animal Care and Use Committee (IACUC). *Esr1*^Cre^ (Lee et al., 2014), *Rpl22*^HA^ (Sanz et al., 2009), *ROSA26*^CAG-Sun1-sfGFP-Myc^ (Mo et al., 2015), and C57Bl6/J wildtype mice were obtained from Jackson labs. Adult male and female mice were used between 8-12 weeks of age. For adult hormone treatment experiments, animals were sacrificed for tissue collection four hours after subcutaneous administration of 5μg estradiol benzoate (EB) (Sigma E8515) suspended in corn oil (Sigma C8267) or vehicle three weeks post-gonadectomy. For the neonatal hormone treatment experiment, animals received a subcutaneous injection of 5μg EB or vehicle on the day of birth and were sacrificed 4 days later.

### Cell lines

Cell lines include mHypoA clu-175 clone (Cedarlane Labs) and MCF7 (ATCC). Cells were maintained in standard DMEM supplemented with 10% FBS and penicillin/streptomycin. Prior to CUT&RUN, MCF7 cells were grown in phenol-red free DMEM media containing 10% charcoal-stripped FBS and penicillin/streptomycin for 48 hours then treated with 20 nM 17-β-estradiol or vehicle (0.002% EtOH) for 45 minutes.

### Adult RNA-seq

Experiments were performed as previously described (Ahrens et al., 2018). Briefly, the BNSTp was microdissected following rapid decapitation of deeply anesthetized adult *Esr1*^Cre/+^; *Rpl22*^HA/+^ mice. Tissue homogenization, immunoprecipitation, and RNA extraction was performed, and libraries were prepared from four biological replicates samples (each consisting of 8-9 pooled animals) using NuGEN Ovation RNA-Seq kits (7102 and 0344). Multiplexed libraries were sequenced with 76bp single end reads on the Illumina NextSeq. Reads were adapter-trimmed and quality-filtered (q>30) (http://hannonlab.cshl.edu/fastx_toolkit/), then mapped to the mm10 reference genome using STAR (Dobin et al., 2013). The number of reads mapping to the exons of each gene was counted with featureCounts (Liao et al., 2014), using the NCBI RefSeq mm10 gene annotation. Differential gene expression analysis was performed using DESeq2 (Love et al., 2014) with the following designs: effect of treatment (design = ~ batch + hormone), effect of sex (design = ~ batch + sex), two-way comparison of treatment and sex (design = ~ batch + hormone_sex), and four-way comparison (design = ~ 0 + hormone_sex). Validation by in-situ hybridization staining and quantification was performed as previously described (Ahrens et al., 2018; Wu and Tollkuhn, 2017). Riboprobe sequences are listed in Table S1.

### ATAC-seq nuclei isolation

Adult *Esr1*^Cre/+^; *ROSA26*^CAG-Sun1-sfGFP-Myc/+^ mice were deeply anesthetized with ketamine/dexmedetomidine. 500-μm sections spanning the BNSTp were collected in an adult mouse brain matrix (Kent Scientific) on ice. The BNSTp was microdissected (4 animals pooled per condition) and collected in 1 ml of cold homogenization buffer (250 mM sucrose, 25 mM KCl, 5 mM MgCl2, 120 mM tricine-KOH, pH 7.8) containing 1 mM DTT, 0.15 mM spermine, 0.5 mM spermidine, and 1X EDTA-free PIC (Sigma Aldrich 11873580001). The tissue was dounce-homogenized 15x in a 1 ml glass tissue grinder (Wheaton) with a loose pestle. 0.3% IGEPAL CA-630 was added, and the suspension was homogenized 5x with a tight pestle. The homogenate was filtered through a 40-μm strainer then centrifuged at 500 × g for 15 min at 4°C. The pellet was resuspended in 0.5 ml homogenization buffer containing 1 mM DTT, 0.15 mM spermine, 0.5 mM spermidine, and 1X EDTA-free PIC. 30,000 GFP+ nuclei were collected into cold ATAC-RSB (10 mM Tris-HCl pH 7.5, 10 mM NaCl, 3 mM MgCl2) using the Sony SH800S Cell Sorter (purity mode) with a 100-μm sorting chip. After sorting, 0.1% Tween-20 was added, and the nuclei were centrifuged at 500 × g for 5 min at 4°C. The nuclei pellet was directly resuspended in transposition reaction mix.

### ATAC-seq library preparation

Tn5 transposition was performed using the OMNI-ATAC protocol (Corces et al., 2017). 2.5 ul of Tn5 enzyme (Illumina 20034197) were used in the transposition reaction. Libraries were prepared with NEBNext High-Fidelity 2X PCR Master Mix (NEB M0541L), following standard protocol. After the initial 5 cycles of amplification, an additional 4 cycles were added, based on qPCR optimization. Following amplification, libraries were size-selected (0.5x - 1.8x) twice with AMPure XP beads (Beckman Coulter A63880) to remove residual primers and large genomic DNA. Individually barcoded libraries were multiplexed and sequenced with paired-end 76bp reads on an Illumina NextSeq, using either the Mid or High Output Kit.

### ATAC-seq data processing

ATAC-seq data were processed using the ENCODE ATAC-seq pipeline (https://github.com/ENCODE-DCC/atac-seq-pipeline) with default parameters. To generate CPM-normalized bigwig tracks, quality-filtered, Tn5-shifted BAM files were converted to CPM-normalized bigwig files using DeepTools bamCoverage (--binSize 1 --normalizeUsing CPM) (Ramírez et al., 2016).

### ATAC-seq differential peak calling

ATAC-seq differential peak calling was performed with DiffBind (Stark et al., 2011). A DiffBind dba object was created from individual replicate BAM and MACS2 narrowPeak files (n = 3 per condition). A count matrix was created with dba.count, and consensus peaks were re-centered to +/−250 bp around the point of highest read density (summits=250). Contrasts between sex and treatment were established (categories = c(DBA_TREATMENT, DBA_CONDITION)), and edgeR (Robinson et al., 2010) was used for differential peak calling. Differential peaks with an FDR<0.05 and log2FC>1 were used for downstream analysis. To call sex-specific, estrogen-responsive peaks shown in Fig. S6, differential peak-calling was performed for 1 condition vs. all other conditions separately, and peaks that were significantly differential across all comparisons were used for heatmap plotting. DeepTools computeMatrix and plotHeatmap were used to plot mean ATAC CPM at EB-open and sex-specific, estrogen-responsive ATAC peaks.

### ATAC-seq data processing

EB-open ATAC peaks and total Veh or EB ATAC peaks (intersected across replicate and sex for each treatment condition) were annotated to NCBI RefSeq mm10 genes using ChIPseeker (Yu et al., 2015a). BETA (basic mode, -d 500000, -c 0.005) (Wang et al., 2013) was used to determine whether EB-open ATAC peaks were significantly overrepresented at differential RNA-seq genes (p<0.01) compared to non-differential, expressed genes. Motif enrichment analysis of EB-open ATAC peaks was performed with AME (Bailey et al., 2009), using the 2020 JASPAR core non-redundant vertebrate database. Motif enrichment analysis was performed using a control file consisting of shuffled primary sequences that preserves the frequency of *k*-mers (--control --shuffle--). FIMO (Grant et al., 2011) was used to determine the percentage of EB-open ATAC peaks containing the enriched motifs shown in Fig. 2E.

### Cell line CUT&RUN

To harvest cells for CUT&RUN, cells were washed twice with Hank’s Buffered Salt Solution (HBSS) and incubated for five minutes with pre-warmed 0.5% Trypsin-EDTA (10X) at 37oC/5% CO_2_. Trypsin was inactivated with DMEM supplemented with 10% FBS and penicillin/streptomycin (mHypoA cells) or phenol-red free DMEM supplemented with 10% charcoal-stripped FBS and penicillin/streptomycin (MCF7 cells). After trypsinizing, cells were centrifuged at 500 × g in a 15ml conical tube and resuspended in fresh media. CUT&RUN was performed as previously described (Skene et al., 2018), with minor modifications. Cells were washed twice in Wash Buffer (20 mM HEPES, pH 7.5, 150 mM NaCl, 0.5 mM spermidine, 1X PIC, 0.02% digitonin). Cell concentration was measured on a Countess II FL Automated Cell Counter (Thermo Fisher). 25,000 cells were used per sample. Cells were bound to 20 ul concanavalin A beads (Bangs Laboratories, <u>BP531</u>), washed 2x in Wash Buffer, and incubated overnight with primary antibody (ERα: Santa Cruz sc-8002, Nfix: Abcam ab101341) diluted 1:100 in Antibody Buffer (Wash Buffer containing 2 mM EDTA). The following day, cells were washed 2x in Wash Buffer, and 700 ng/ml protein A-MNase (pA-MNase, prepared in-house) was added. After 1 hr incubation at 4oC, cells were washed 2x in Wash Buffer and placed in a metal heat block on ice. pA-MNase digestion was initiated with 2 mM CaCl2. After 90 min, digestion was stopped by mixing 1:1 with 2X Stop Buffer (340 mM NaCl, 20 mM EDTA, 4 mM EGTA, 50 ug/ml RNase A, 50 ug/ml glycogen, 0.02% digitonin). Digested fragments were released by incubating at 37°C for 10 min, followed by centrifuging at 16,000 × g for 5 min at 4°C. DNA was purified from the supernatant by phenol-chloroform extraction, as previously described (Skene et al., 2018).

### Brain nuclei CUT&RUN

Nuclei were isolated from microdissected POA, BNSTp, and MeAp from gonadectomized C57Bl6/J mice, following anatomic designations (Paxinos and Franklin, 2019) (Fig. 1A), as described previously (Mo et al., 2015). Following tissue douncing, brain homogenate was mixed with a 50% OptiPrep solution and underlaid with 4.8 ml of 30% then 40% OptiPrep solutions, in 38.5 ml Ultra-clear tubes (Beckman-Coulter C14292). Ultracentrifugation was performed with a Beckman SW-28 swinging bucket rotor at 9200 RPM for 18 minutes at 4°C. Following ultracentrifugation, ~1.5 ml of nuclei suspension was collected from the 30/40% OptiPrep interface by direct tube puncture with a 3 ml syringe connected to an 18-gauge needle. Nuclei concentration was measured on a Countess II FL Automated Cell Counter. For ERα CUT&RUN, 400,000 nuclei were isolated from BNST, MPOA, and MeA of 5 animals. For Nfix CUT&RUN, 200,000 nuclei were isolated from BNSTp of 5 animals. 400,000 cortical nuclei were used for the CUT&RUN IgG control. Prior to bead binding, 0.4% IGEPAL CA-630 was added to the nuclei suspension to increase affinity for concanavalin A magnetic beads. All subsequent steps were performed as described above, with a modified Wash Buffer (20 mM HEPES, pH 7.5, 150 mM NaCl, 0.1% BSA, 0.5 mM spermidine, 1X PIC) and an ERα antibody recognizing the mouse epitope (Millipore Sigma 06-935).

### CUT&RUN library preparation

Cell line CUT&RUN libraries were prepared using the SMARTer ThruPLEX DNA-seq Kit (Takara Bio R400676), with the following PCR conditions: 72°C for 3 min, 85°C for 2 min, 98°C for 2 min, (98°C for 20 sec, 67°C for 20 sec, 72°C for 30 sec) × 4 cycles, (98°C for 20 sec, 72°C for 15 sec) × 14 cycles (MCF7) or 10 cycles (mHypoA). Brain CUT&RUN libraries were prepared using the same kit with 10 PCR cycles. All samples were size-selected with AMPure XP beads (0.5x - 1.7x) to remove residual adapters and large genomic DNA. Individually barcoded libraries were multiplexed and sequenced with paired-end 76bp reads on an Illumina NextSeq, using either the Mid or High Output Kit. For the mHypoA experiment, samples were sequenced with paired-end 25bp reads on an Illumina MiSeq.

### CUT&RUN data processing

Paired-end reads were trimmed to remove Illumina adapters and low-quality basecalls (cutadapt -q 30) (Martin, 2011). Trimmed reads were aligned to mm10 using Bowtie2 (Langmead and Salzberg, 2012) with the following flags: --dovetail --very-sensitive-local --no-unal --no-mixed --no-discordant --phred33. Duplicate reads were removed using Picard (http://broadinstitute.github.io/picard/) MarkDuplicates (REMOVE_DUPLICATES=true). Reads were filtered by mapping quality (samtools view -q 40) and fragment length (deepTools alignmentSieve -- maxFragmentLength 120). Reads aligning to the mitochondrial chromosome and incomplete assemblies were also removed using samtools. After filtering, peaks were called on individual replicate BAM files using MACS2 callpeak (--min-length 25 -q 0.01) (Zhang et al., 2008). To identify total Nfix peaks, MACS2 callpeak was performed on BAM files merged across biological replicates and subsequently intersected across treatment and sex. Sample peaks that overlapped peaks called in the IgG control were removed using bedtools intersect (-v) (Quinlan and Hall, 2010) prior to downstream analysis.

### CUT&RUN differential peak calling

CUT&RUN differential peak calling was performed with DiffBind. A count matrix was created from individual replicate BAM and MACS2 narrowpeak files (n = 2 per condition). Consensus peaks were re-centered to +/−100 bp around the point of highest read density (summits=100). Differential peak calling was performed, as described above, using edgeR with block set to DBA_REPLICATE. Differential ERα peaks with an padj<0.1 were used for downstream analysis. For Nfix, differential peaks with an padj<0.1 and abs(log2FC) > 1 were used for downstream analysis.

### CUT&RUN data analysis

EB-induced ERα peaks were annotated to NCBI RefSeq mm10 genes using ChIPseeker, as described above. DeepTools plotHeatmap was used to plot ERα CUT&RUN (Fig. 1B) and ATAC coverage (Fig. 2H), representing CPM-normalized bigwig files pooled across replicate and sex per condition, at EB-induced ERα peaks. Heatmaps of individual ERα CUT&RUN replicates are shown in Fig. S2. CUT&RUNTools (Zhu et al., 2019) was used to plot ERα CUT&RUN fragment ends surrounding ESR1 motifs (JASPAR MA0112.3) within EB-induced ERα peaks as well as Nfix CUT&RUN fragment ends surrounding NFIX dimer motifs (JASPAR MA1528.1). The BETA tool was used to assess statistical association between ERα peaks and differential RNA-seq genes, as done for EB-open ATAC-seq peaks. Motif enrichment analysis of ERα peaks was performed with AME using both the 2020 JASPAR core non-redundant vertebrate database (Fig. S2) and the 2020 JASPAR vertebrata CORE PWM matrix clusters (http://jaspar2020.genereg.net/matrix-clusters/) (Fig. 3), which had lower motif redundancy. The following 7 ERα ChIP-seq files were lifted over to mm10 using UCSC liftOver and intersected with EB-induced ERα peaks to identify brain-specific and shared (≥4 intersections) ERα binding sites: uterus [intersection of GEO:GSE36455 (uterus 1) (Hewitt et al., 2012) & GEO:GSE49993 (uterus 2) (Gertz et al., 2013)], liver [intersection of GEO:GSE49993 (liver 1) (Gertz et al., 2013) & GEO:GSE52351(liver 2) [(Gordon et al., 2014)], aorta [(Gordon et al., 2014), GEO: GSE52351], efferent ductules [(Yao et al., 2017), Supplementary information], and mammary gland [(Palaniappan et al., 2019), GEO: GSE130032]. The same approach was used to identify ERα ChIP-seq peaks specific to each dataset. Brain ERα peaks overlapping at least 4 of the external datasets (counted with bedtools intersect -C) were classified as “shared”. DeepTools plotProfile was used to plot BNSTp vehicle ATAC coverage at tissue-specific ERα peaks. Mouse liver ATAC-seq data (Cusanovich et al., 2018) were accessed from the mouse chromatin atlas (https://shendurelab.github.io/mouseatac/data/), normalized using DeepTools bamCoverage, lifted over to mm10 using CrossMap (Zhao et al., 2014), and plotted in the same way as BNSTp vehicle ATAC data. Genomic annotation of brain-specific and shared ERα peaks was performed with ChIPseeker. clusterProfiler (Yu et al., 2012) was used to identify associations between brain-specific and shared ERα peak-annotated genes and Gene Ontology (GO) Biological Process terms (enrichGO, ont=“BP”, padj<0.1). For Disease Ontology (DO) associations, brain-specific ERα peak-associated gene symbols were converted from mouse to human using bioMart (Durinck et al., 2009) then analyzed with DOSE (Yu et al., 2015b) (enrichDO, padj<0.1). Log-odds ESR1 motif scores in brain-specific and shared ERα peaks were calculated with FIMO, using default parameters.

MCF7 ERα CUT&RUN data were compared to MCF7 ERα ChIP-seq data from [(Franco et al., 2015); GEO: GSE59530]. Single-end ChIP-seq fastq files for 2 vehicle-treated and 2 17β-estradiol (E2)-treated IP and input samples were accessed from Sequence Read Archive and processed identically as ERα CUT&RUN data, with the exception of fragment size filtering. Differential ERα ChIP-seq peak calling was performed using DiffBind DESeq2 (padj<0.01). DeepTools was used to plot CPM-normalized ERα CUT&RUN signal at E2-induced ERα ChIP-seq binding sites. DREME (Bailey, 2011) and AME were used to compare *de novo* and enriched motifs between E2-induced MCF7 ERα CUT&RUN and ChIP-seq peaks.

Bedtools intersect (-wa) was used to identify overlapping regions (≥1bp) between ERα peaks, Nfix non-differential peaks (total peaks minus differential peaks), and differential Nfix peaks. Mean vehicle and EB ATAC CPM at ERα+Nfix pre-bound (ERα + non-differential Nfix) and ERα+Nfix recruited (ERα + differential Nfix) sites was computed using DeepTools multiBigwigSummary (BED-file mode). Log-odds ESR1 motif scores in ERα+Nfix pre-bound and ERα+Nfix recruited sites were determined with FIMO, as done previously.

### Pup RNA-seq

Female *Esr1*^Cre/+^; *ROSA26*^CAG-Sun1-sfGFP-Myc/+^ mice were injected subcutaneously with 5μg EB or vehicle on P0. Four days later, animals were rapidly decapitated, and 400-μm sections were collected in ice-cold homogenization buffer (250 mM sucrose, 25 mM KCl, 5 mM MgCl2, 120 mM tricine-KOH, pH 7.8) using a microtome (Thermo Scientific Microm HM 650V). The BNST was microdissected (4 animals pooled per condition) and collected in 1 ml of cold homogenization buffer containing 1 mM DTT, 0.15 mM spermine, 0.5 mM spermidine, and 1X EDTA-free PIC, and 0.4 U/ml RNAseOUT (Thermo Fisher, 10777019). Nuclei isolation was performed as done for adult ATAC-seq, with the addition of 0.2 U/ml RNAseOUT to the resuspension buffer. 12,000 GFP+ nuclei were collected into cold Buffer RLT Plus supplemented 1:100 with β-mercaptoethanol (Qiagen, 74034) using the Sony SH800S Cell Sorter (purity mode) with a 100-μm sorting chip (same gating strategy used for adult ATAC-seq). Nuclei were stored at −80°C until all replicates were collected. Nuclei samples for all replicates were thawed on ice, and RNA was isolated using the Qiagen RNeasy Plus Micro Kit. Strand-specific RNA-seq libraries were prepared using the Ovation SoLo RNA-seq system (Tecan Genomics, 0501-32), following manufacturer’s guidelines. Individually barcoded libraries were multiplexed and sequenced with single-end 76bp reads on an Illumina NextSeq, using the Mid Output Kit. Reads were trimmed to remove Illumina adapters and low-quality basecalls (cutadapt -q 30), then mapped to the mm10 reference genome using STAR. Technical duplicate reads (identical start and end positions with same strand orientation and identical molecular identifier) were removed using the nudup.py python package (https://github.com/tecangenomics/nudup). The number of reads mapping to each gene (including introns) on each strand (-s 1) was calculated with featureCounts (Liao et al., 2014), using the mm10.refGene.gtf file. Differential gene expression analysis was performed using DESeq2 (design = ~ treatment) after prefiltering genes by expression (rowMeans>=5).

### Single-nucleus and -cell RNA-seq analysis

Mouse BNST snRNA-seq data containing 76,693 neurons across 7 adult female and 8 adult male biological replicates (Welch et al., 2019) were accessed from GEO:GSE126836 and loaded into a Seurat object (Stuart et al., 2019). Mouse MPOA/BNST scRNA-seq data containing 31,299 cells across 3 adult female and 3 adult male biological replicates (Moffitt et al., 2018) were also accessed from GEO:GSE113576 and loaded into a Seurat object. Cluster identity, replicate, and sex were added as metadata features to each Seurat object. Pseudo-bulk RNA-seq analysis was performed to identify sex differences in gene expression in the BNST snRNA-seq dataset. Briefly, the Seurat object was converted to a SingleCellExperiment object (as.SingleCellExperiment). Genes were filtered by expression (genes with >1 count in ≥5 nuclei), and NCBI predicted genes were removed. For each cluster, nuclei annotated to the cluster were subsetted from the main Seurat object. Biological replicates containing 20 or fewer nuclei in the cluster were excluded. Gene counts for each biological replicate with >20 nuclei in the cluster were aggregated. Differential gene expression analysis across sex within each cluster was performed on the filtered, aggregated count matrix using DESeq2 (design = ~ sex) with alpha = 0.1. The BNSTp_Cplx3 cluster was excluded, as none of the replicates in this cluster contained over 20 nuclei. Clusters containing ≥25% nuclei with ≥1 *Esr1* counts in the main Seurat object were classified as *Esr1*-expressing/*Esr1*+ (BNST_Avp, BNSTp_Tac2, BNSTa_Esr1, BNSTpr_Esr2, BNSTa_Synpo2, BNSTp_Bnc2, BNSTpr_St18, BNSTa_Th, BNSTp_Epsti1, BNST_C1ql3, BNSTa_Zeb2, BNSTal_Chat, BNSTa_Gli3, BNSTp_Nxph2, BNSTa_Lrrc9, BNSTp_Haus4).

To visualize BNSTp *Esr1*-expressing nuclei, these clusters were subsetted from the main Seurat object. Gene counts were log-normalized, and the top 2000 variable features were identified using FindVariableFeatures (selection.method = ‘vst’). Gene counts were scaled, and linear dimensionality reduction was performed by principal component analysis (runPCA, npcs = 10). BNSTp *Esr1*-expressing clusters were visualized with UMAP, using the same number of dimensions as PCA (runUMAP, dims = 10). BETA (basic mode, -d 500000, -c 0.005) was used to infer direct ERα target genes among sexually dimorphic genes (DESeq2, padj<0.1) in each cluster. To generate the heatmap in Fig. 4D, pseudo-bulk counts for each biological replicate were normalized and transformed with variance stabilizing transformation (DESeq2 ‘vst’), subsetted for male-biased ERα target genes in each cluster, and z-scaled across replicates. ERα CUT&RUN CPM at the single nearest ERα peak associated with each male-biased gene was plotted using DeepTools. The order of both snRNA-seq and CUT&RUN heatmaps was sorted by EB ERα CUT&RUN CPM.

To identify marker genes enriched in the BNSTpr St18 cluster relative to the other 6 *Esr1*+ BNSTp/pr clusters (Fig. S9), differential gene expression analysis was performed on the filtered, aggregated count matrix using DESeq2 with design = ~ cluster_id (betaPrior = TRUE), alpha = 0.01, lfc-Threshold = 2, altHypothesis = “greater”.

*Esr1*-expressing BNSTp and pr clusters were integrated with MPOA/BNST *Esr1*-expressing clusters (e3: Cartpt_Isl1, i18: Gal_Tac2, i20: Gal_Moxd1, i28: Gaba_Six6, i29: Gaba_Igsf1, i38: Kiss1_Th) using Seurat. ‘Anchors’ were identified between cells from the two unscaled datasets, using FindIntegrationAnchors. An integrated expression matrix was generated using IntegrateData (dims = 1:10). The resulting integrated matrix was used for downstream PCA and UMAP visualization (dims = 1:10).

---

**Fig. S1.**
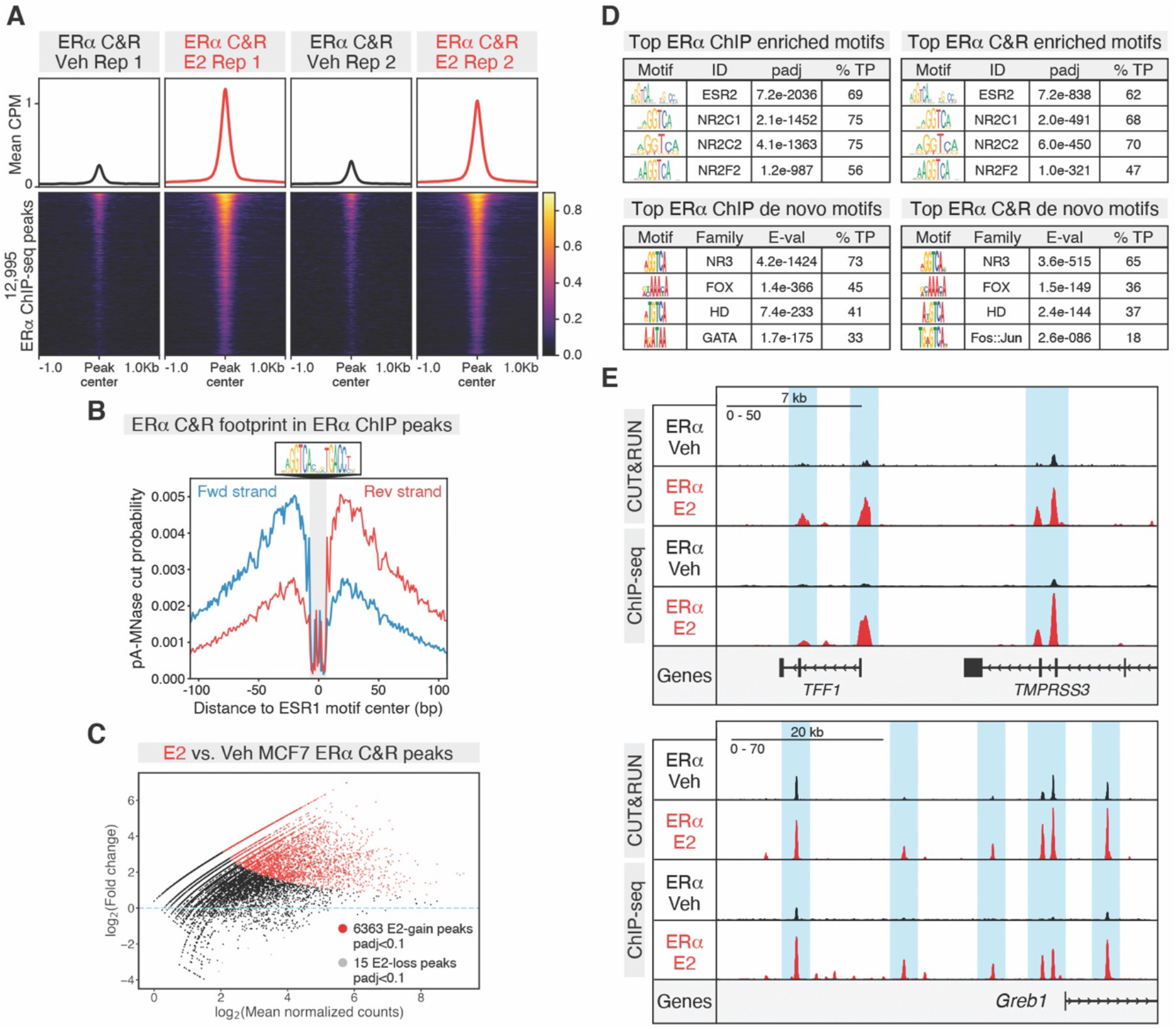
CUT&RUN enables low input profiling of ERα binding in MCF7 breast cancer cells. **(A)** Heatmap of mean Veh and E2 MCF7 ERα CUT&RUN CPM (n=2) +/−1Kb around center of 17β-estradiol (E2)-induced ERα ChIP-seq peaks (DiffBind DESeq2, padj<0.01). Heatmap sorted by mean CPM across the rows. **(B)** pA-MNase-cut footprint (CUT&RUNTools) around ESR1 motif sites in 6,363 E2-induced ERα CUT&RUN peaks (DiffBind DESeq2, padj<0.1). **(C)** MA plot of E2 vs. Veh differential ERα CUT&RUN peaks. Red dots=E2-induced sites (padj<0.1); grey dots=Veh-induced sites (padj<0.1). **(D)** (Top) Top enriched motifs (AME) and (bottom) *de novo* motifs in E2-induced ERα ChIP-seq peaks (left) and ERα CUT&RUN peaks (right). % TP=% of peaks called as positive for the indicated motif. *De novo* motifs were classified into motif families using TomTom. **(E)** Example ERα CUT&RUN and ChIP-seq peaks at canonical MCF7 loci *TFF1* and *GREB1*.

**Fig. S2.**
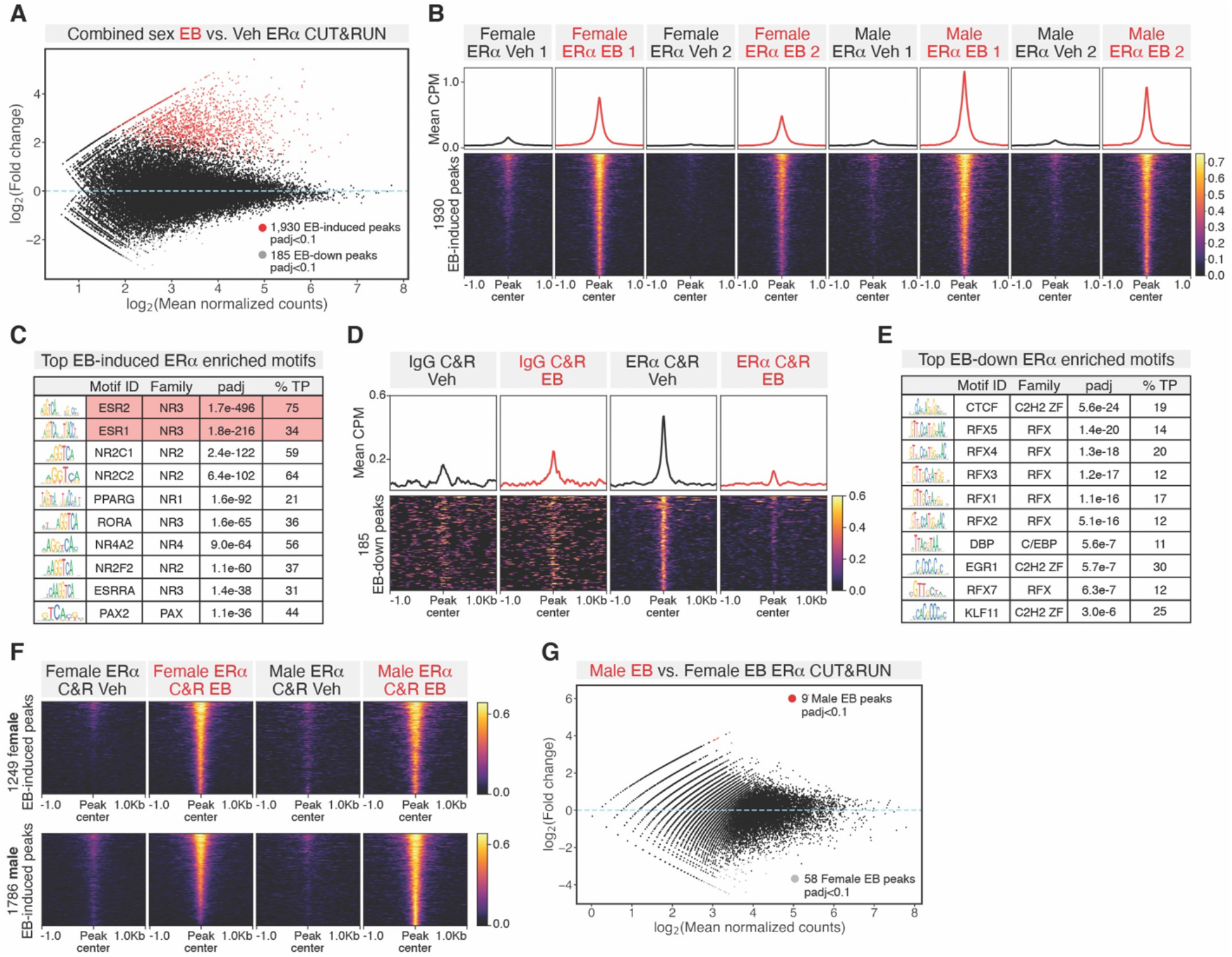
Additional analysis of brain ERα CUT&RUN experiment. **(A)** MA plot of EB vs. Veh differential ERα CUT&RUN peaks in the brain (DiffBind edgeR, padj<0.1). Red dots=EB-induced sites; grey dots=EB-down sites. **(B)** Heatmap of mean ERα CUT&RUN CPM +/−1Kb around 1,930 EB-induced peaks for individual female and male replicates (n=2 per condition, see also Fig. 1B). Heatmap sorted by mean CPM across the rows. **(C)** Top motifs enriched (AME) in ERα peaks from the JASPAR 2020 non-redundant vertebrate core database, without motif clustering (see also Fig. 3A). % TP=% of peaks called as positive for the indicated motif. **(D**) Heatmap of mean IgG and ERα CUT&RUN CPM +/−1Kb around 185 EB-down (vehicle) ERα CUT&RUN peaks. Heatmap sorted by ERα Veh CUT&RUN CPM. **(E)** Top motifs enriched (AME) in EB-down peaks. **(F)** Heatmap of mean ERα CUT&RUN CPM +/−1Kb around (top) 1,249 EB-induced peaks detected in females and (bottom) 1,786 EB-induced peaks detected in males (DiffBind edgeR, padj<0.1). Both heatmaps sorted by mean CPM across the rows. At peaks called for each sex, there is EB-induced ERα recruitment detected in the opposite sex, demonstrating lack of sex-specific ERα recruitment. **(G)** MA plot of Male EB vs. Female EB ERα CUT&RUN peaks (DiffBind edgeR, padj<0.1). Red dots=Male EB-biased sites; grey dots=Female EB-biased sites.

**Fig. S3.**
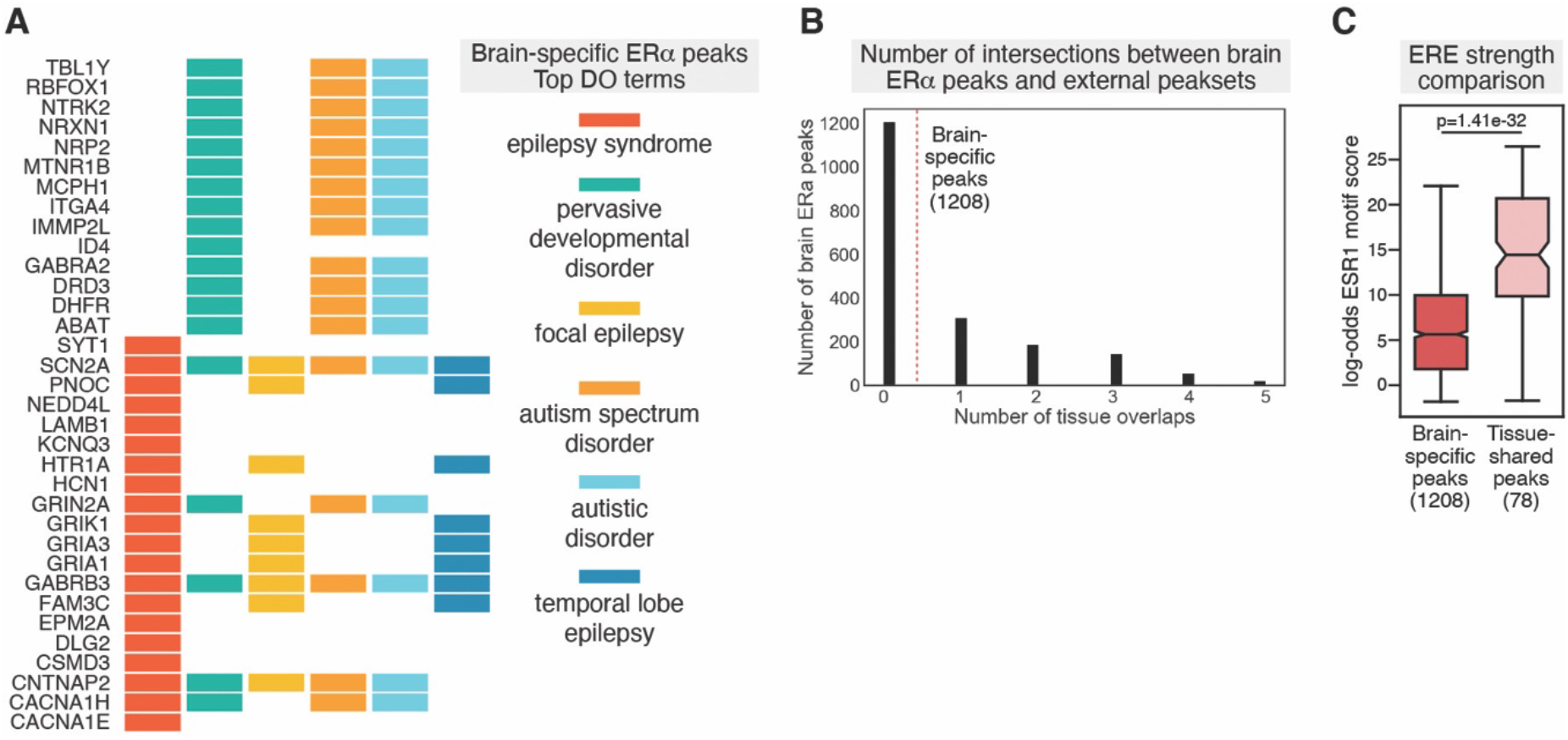
Additional analysis of brain-specific vs. shared ERα binding sites. **(A)** Genes annotated to brain-specific ERα peaks (ChIPSeeker) in the top enriched Disease Ontology (DO) terms. Genes are colored by term. **(B)** Number of intersections between brain ERα CUT&RUN peaks and 7 external ERα ChIP-seq peaksets: intersected peaks of uterus 1 and uterus 2, intersected peaks of liver 1 and liver 2, aorta, efferent ductules, and mammary gland. **(C)** Log-odds ESR1 motif score (FIMO, MA0112.3) for brain-specific (1208) and tissue-shared (78) ERα peaks. p-value from Mann-Whitney *U* test.

**Fig. S4.**
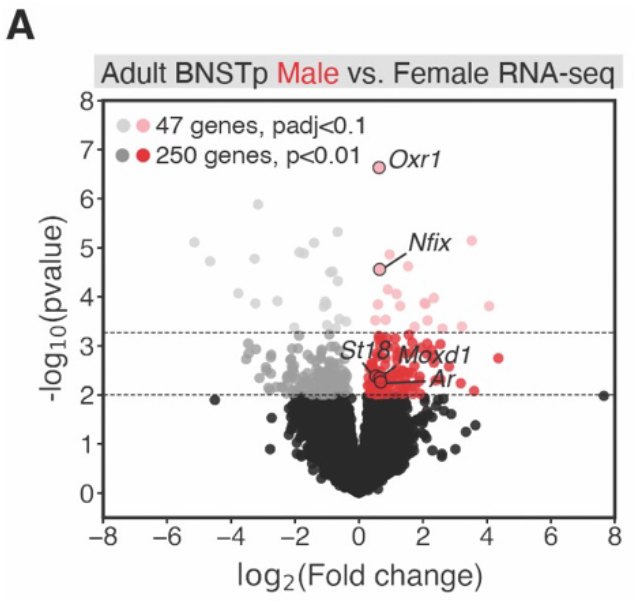
Sexually dimorphic genes in adult BNSTp RNA-seq. **(A)** Volcano plot of sexually dimorphic genes in BNSTp ERα neurons; light grey and red dots (47 genes, DESeq2, padj<0.1), dark grey and red dots (250 genes, DESeq2, p<0.01). Bulk BNSTp ERα+ RNA-seq captures male-biased expression of BNSTpr St18 cluster markers *Nfix*, *Moxd1*, and *St18*.

**Fig. S5.**
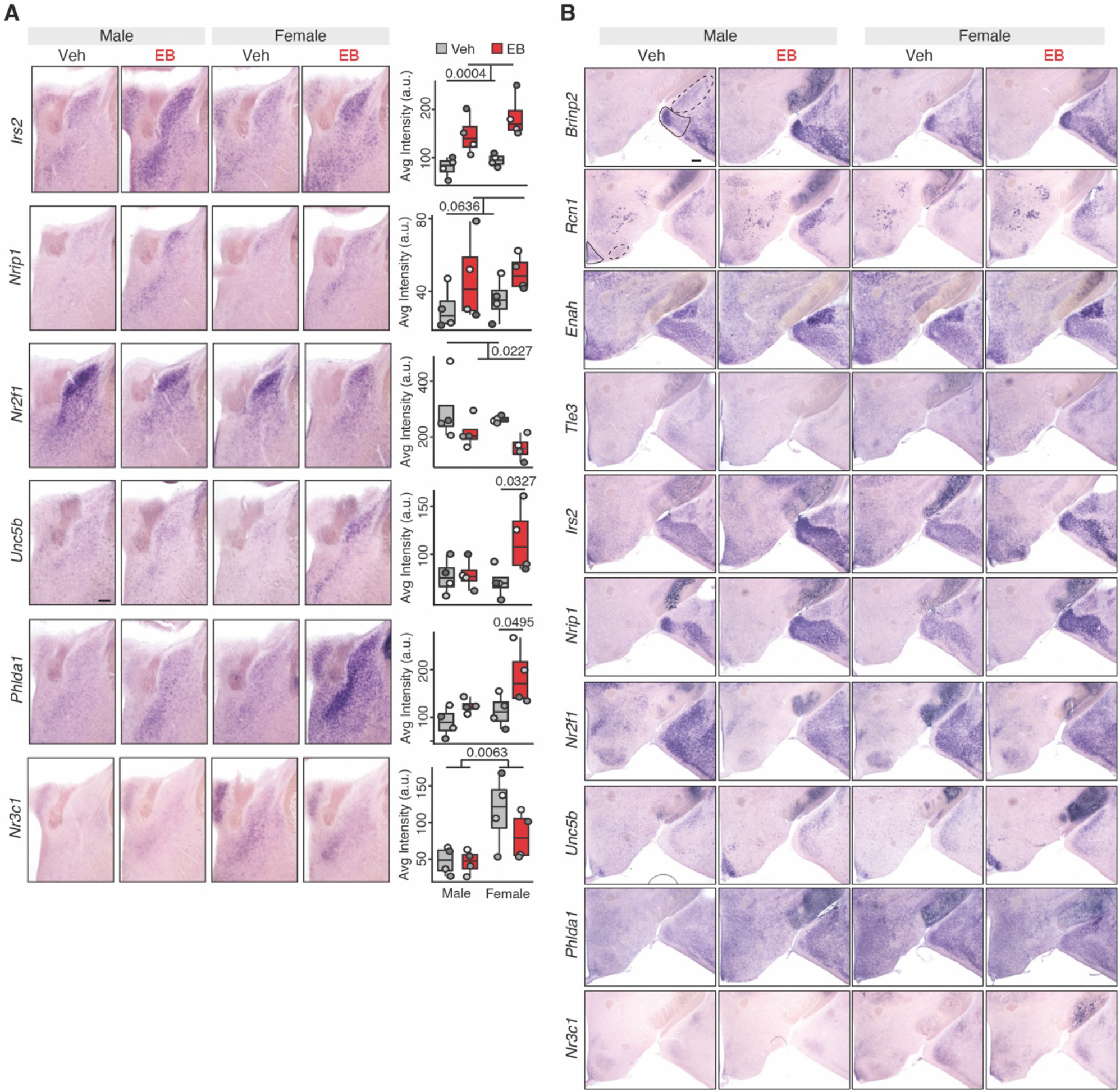
Estrogen regulation of gene expression in additional ERα+ brain regions. **(A)** ISH of select genes in the BNSTp induced by EB in females only (paired t-test *Unc5b* p=0.0327, *Phlda1* p=0.0495); downregulated by EB in both sexes (*Nr2f1* 2-way ANOVA effect of treatment p=0.0227); more highly expressed in females treated with EB or vehicle compared to males (*Nr3c1* 2-way ANOVA effect of sex p=0.0063); or not significantly differentially expressed (*Nrip1* 2-way ANOVA effect of treatment p=0.0636) n=4, scale=200um. **(B)** Representative ISH images of select genes demonstrate that estrogen-induced genes in the BNSTp (see also Fig. 1C) are differentially regulated in other regions. Outlined ROI: posterodorsal medial amygdala (row 1 dotted line), posteroventral medial amygdala (row 1 solid line), ventrolateral ventromedial hypothalamus (row 2 dotted line), and arcuate (row 2 solid line).

**Fig. S6.**
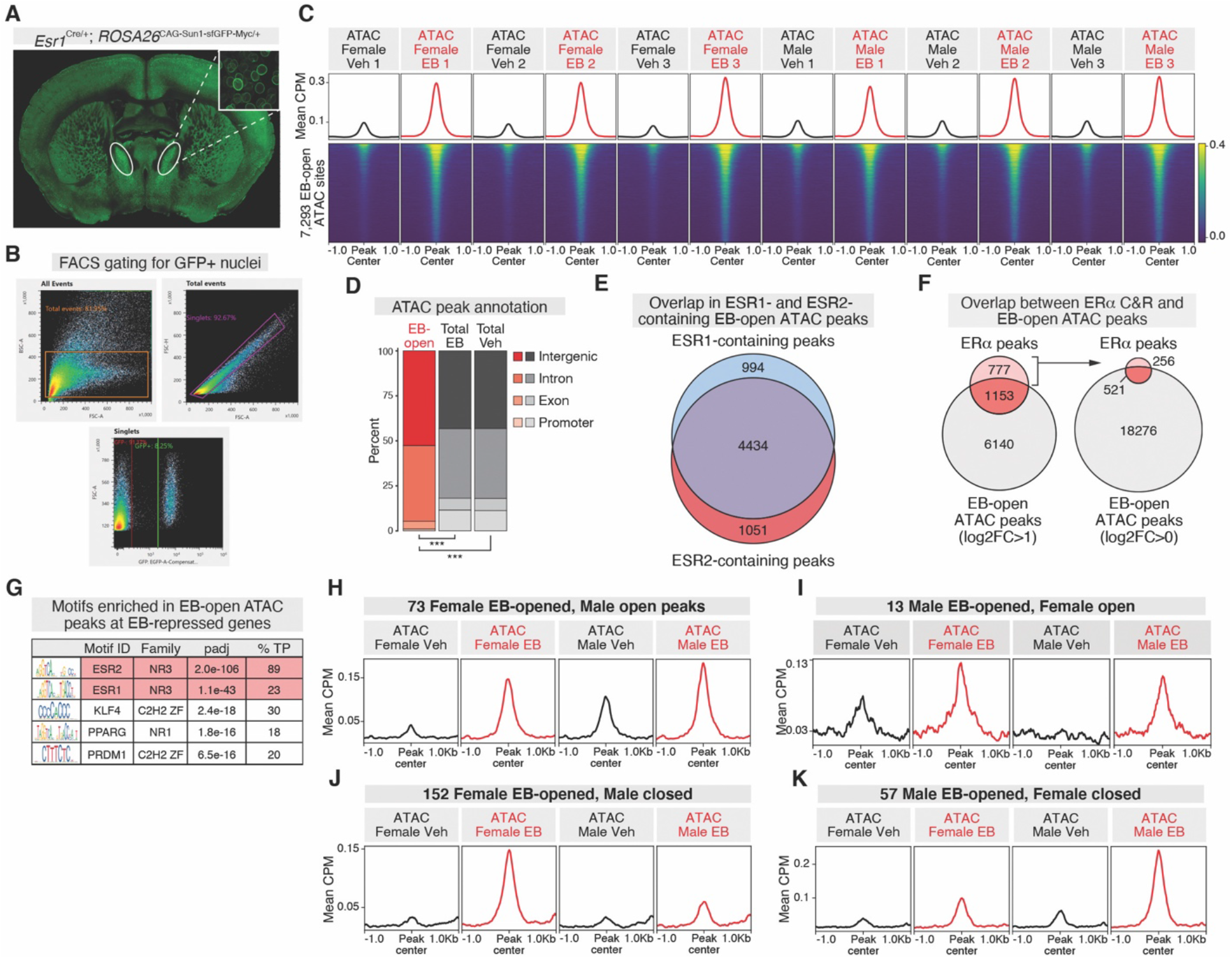
BNSTp ATAC-seq quality control, genomic annotation, and motif enrichment. **(A)** Coronal section of adult C57/Bl6 male *Esr1*^Cre/+^; *ROSA26*^CAG-Sun1-sfGFP-Myc/+^ animal. White circle indicates BNSTp. Inset shows Sun1-GFP signal at nuclear membrane. **(B)** Fluorescence-activated cell sorting (FACS) gating strategy for isolating BNSTp GFP+ nuclei. **(C)** Heatmap of mean ATAC-seq CPM +/−1Kb around 7,293 EB-open ATAC-seq peaks (DiffBind edgeR, log2FC>1, padj<0.05) for individual female and male replicates (n=3 per condition). (**D**) Proportion of EB-open, total vehicle, and total EB ATAC peaks at promoters (+/−1Kb around TSS), exons, introns, and intergenic regions. EB-open ATAC peaks have a significantly lower proportion of’peaks annotated to gene promoters than total vehicle (11% vs 1%, Fisher’s Exact Test, p=4.6×10^−260^) and total EB (11% vs 1%, Fisher’s Exact Test, p=4.3×10^−267^) peaks. ***p<0.001. **(E**) Intersection of EB-open ATAC-seq peaks containing an ESR1 or ESR2 motif. The majority of peaks (4434/6479) that contain either motif are the same. **(F)** (Left) Overlap between ERα CUT&RUN and EB-open ATAC-peaks (log2FC>1, padj<0.05). The majority of ERα peaks (521/777) not overlapping log2FC>1 EB-open ATAC-peaks overlap EB-open ATAC peaks called at log2FC>0. **(G)** Top motifs enriched (AME) in EB-open ATAC-peaks associated with EB-repressed genes (see also Fig. 2F). **(H-K)** Identification of a small number of ATAC peaks that are: **(H)** EB-opened in females and constitutively open in males (78 peaks), **(I)** EB-opened in males and constitutively open in females (13 peaks), **(J)** EB-opened in females and constitutively closed in males (152 peaks), and **(K)** EB-opened in males and constitutively closed in females (57 peaks). padj<0.05 across all comparisons.

**Fig. S7.**
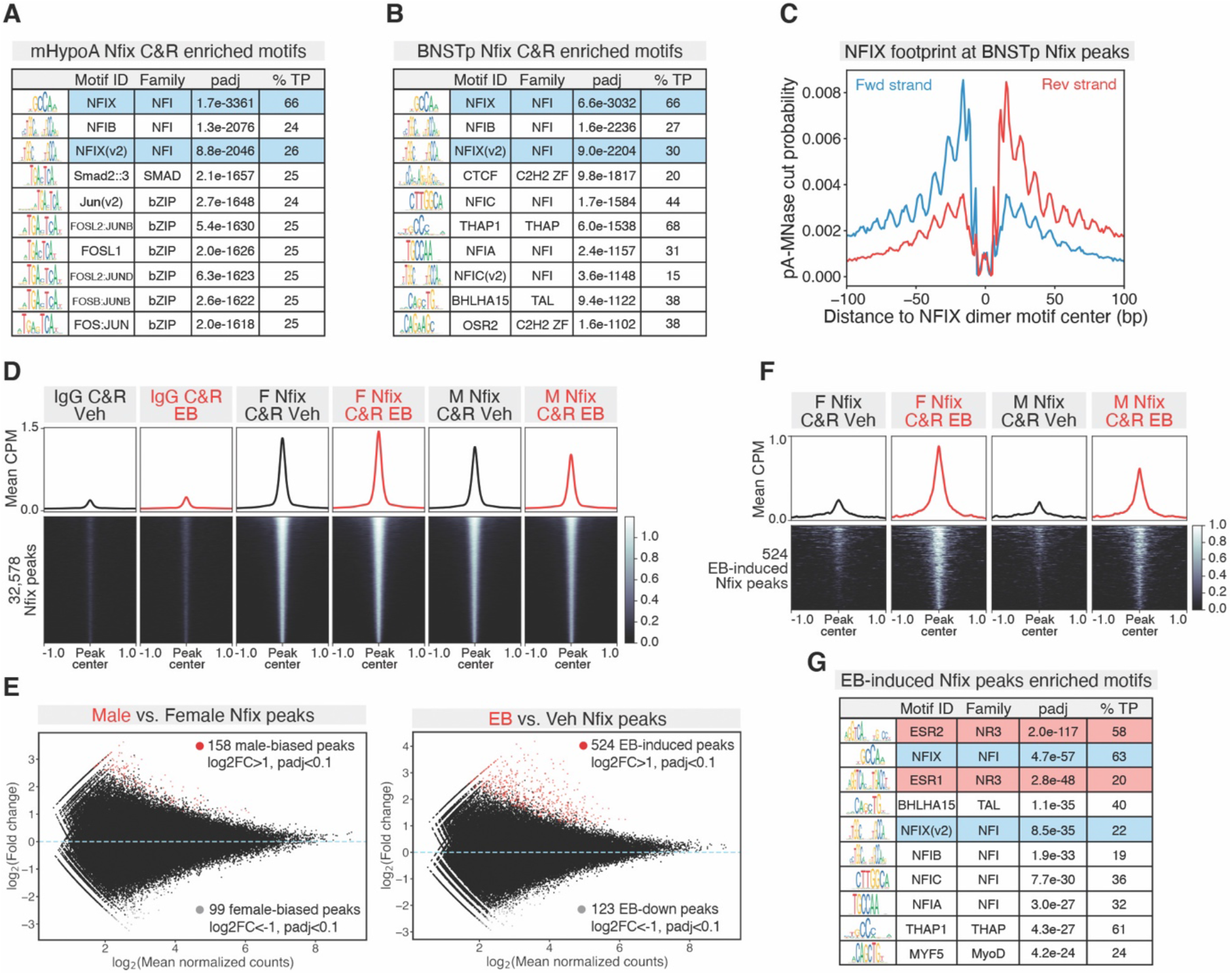
Nfix CUT&RUN in mHypoA cells and BNSTp. **(A)** Top motifs enriched (AME) in 30,825 mHypoA cell Nfix CUT&RUN peaks (MACS2 callpeak, q<0.01). %TP=% of peaks called as positive for the indicated motif. **(B)** Top motifs enriched (AME) in 32,578 total BNSTp Nfix CUT&RUN peaks (MACS2 callpeak, q<0.01; peaks intersected across treatment and sex). %TP=% of peaks called as positive for the indicated motif. **(C)** pA-MNase-cut footprint (CUT&RUNTools) at Nfix dimer motif in total BNSTp Nfix peaks. **(D)** Heatmap of mean IgG and Nfix CUT&RUN CPM (n=2 per condition) +/−1Kb around 32,578 total Nfix peaks. **(E)** MA plot of (left) Male vs. Female and (right) EB vs. Veh differential BNSTp Nfix CUT&RUN peaks (DiffBind edgeR). Red dots=Male-biased (left) and EB-induced (right) sites (log2FC>1, padj<0.1); grey dots=Female-biased (left) and EB-down (right) sites (log2FC<-1, padj<0.1). **(F)** Heatmap of mean Nfix CUT&RUN CPM +/−1Kb around 524 EB-induced peaks in both sexes. Heatmaps sorted by mean CUT&RUN CPM across rows. **(G)** Top motifs enriched (AME) in 524 EB-induced Nfix CUT&RUN peaks.

**Fig. S8.**
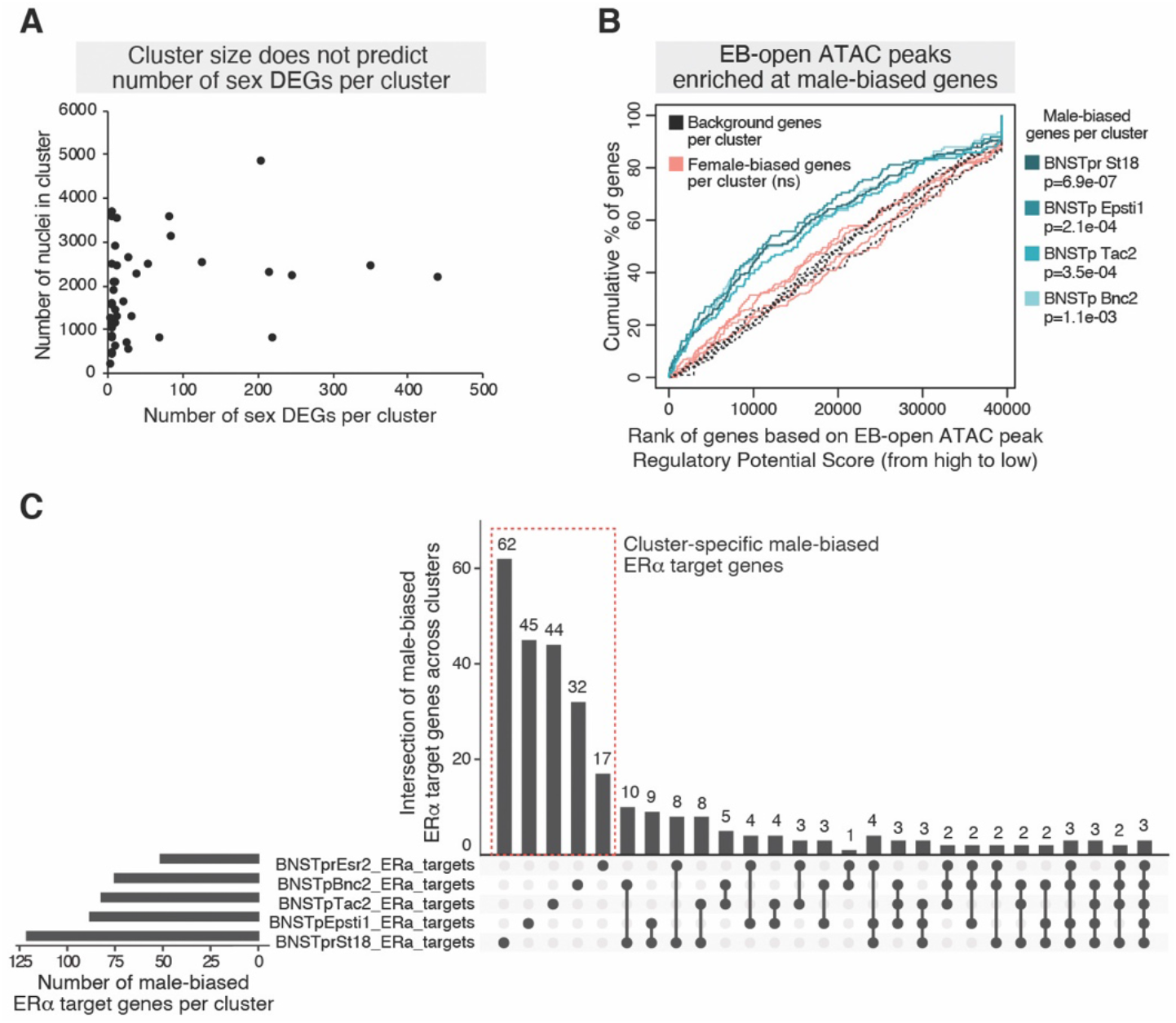
Additional analysis of BNST snRNA-seq dataset. **(A)** Number of nuclei in each BNST snRNA-seq cluster vs. number of sex DEGs (DESeq2, padj<0.1) detected in the cluster. No correlation is detected between cluster size and number of sex DEGs (R^2^=0.068). **(B)** BETA analysis cumulative distribution curve of regulatory potential scores for EB-open ATAC peaks associated with male-biased genes in 4 of the 7 BNSTp *Esr1*+ clusters. p-values from K-S test, ns=not significant. **(C)** Upset plot showing the intersection of male-biased ERα target genes between the 5 BNSTp *Esr1*+ clusters, in which significant over-representation of ERα at male-biased genes was detected. The majority of male-biased ERα target genes are detected in individual clusters (red dotted box).

**Fig. S9.**
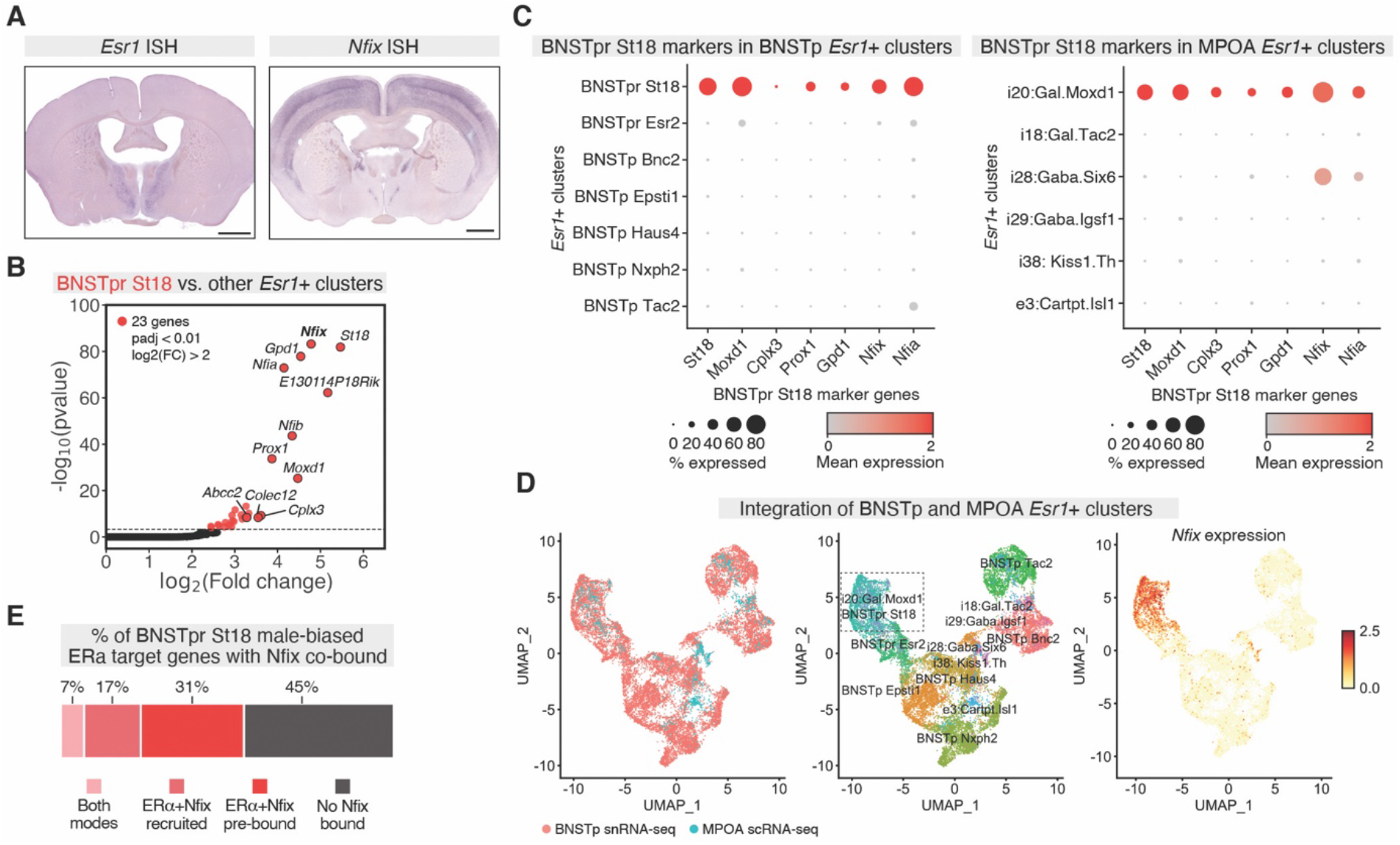
BNSTpr St18 neurons co-cluster with an inhibitory MPOA neuron cluster and co-express *Nfix*. **(A)** While more widely expressed in the cortex, *Nfix* is present in a subset of the BNST and MPOA that co-expresses *Esr1*. ISH of adult veh-treated castrated male (*Esr1*) and adult male (*Nfix*). Scale=1mm. **(B)** Volcano plot of pseudo-bulk differential gene expression analysis between BNSTpr St18 cluster and the 6 other BNSTp *Esr1*+ clusters combined (BNSTp Bnc2, BNSTp Epsti1, BNSTp Haus4, BNSTp Nxph2, BNSTp Tac2, BNSTpr Esr2). Red dots=BNSTpr St18-enriched genes (DESeq2 padj<0.01, log2FC>2). **(C)** Mean, normalized expression of top BNSTpr St18 marker genes in (left) BNSTp *Esr1*+ clusters and (right) MPOA *Esr1*+ clusters. Similar marker gene enrichment detected in BNSTpr St18 and i20:Gal.Moxd1 neurons. **(D)** Integrated clustering of BNSTp *Esr1*+ clusters (BNSTp Bnc2, BNSTp Epsti1, BNSTp Haus4, BNSTp Nxph2, BNSTp Tac2, BNSTpr Esr2, BNSTpr St18) (blue) and MPOA *Esr1*+ clusters (e3:Cartpt.Isl1, i18:Gal.Tac2, i20:Gal.Moxd1, i28:Gaba.Six6, i29:Gaba.Igsf1, i38:Kiss1.Th) (pink) reveal co-clustering of BNSTpr St18 and i20:Gal.Moxd1 neuron clusters and mutual expression of *Nfix*. **(E)** Proportion of BNSTpr St18 male-biased ERα target genes associated with an ERα+Nfix pre-bound site (dark red), ERα+Nfix recruited site (medium red), or both types of sites (light red). Proportion of ERα target genes without Nfix co-bound colored in grey.

**Fig. S10.**
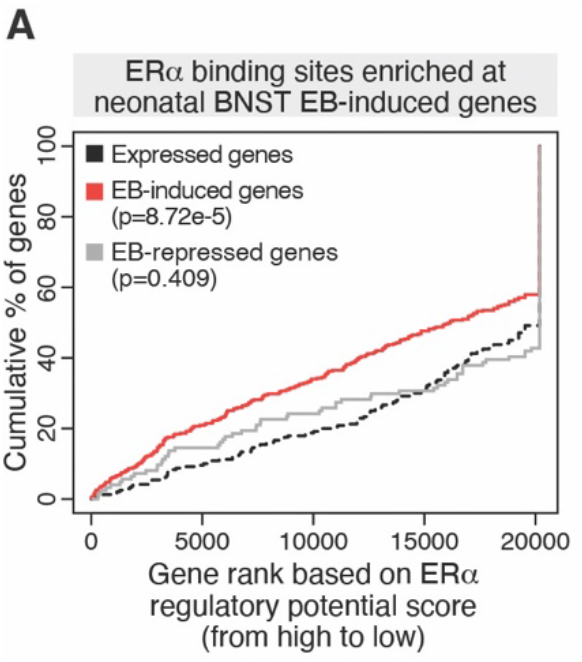
ERα binding enrichment at neonatal EB-induced genes in the BNST. **(A)** BETA analysis cumulative distribution curve of regulatory potential scores for brain ERα CUT&RUN peaks associated with neonatal (P4) BNST EB-regulated (DESeq2, padj<0.05) and non-differential, expressed genes. p-values from K-S test.

**Table S1.** In situ hybridization riboprobes.

| **Gene** | **RefSeqID** | **Range** |
| --- | --- | --- |
| Brinp2 | NM_207583.2 | 2221-3115 |
| Esr1 | NM_001302532.1 | 1263-2243 |
| Enah | NM_001083120.2 | 2597-3067 |
| Irs2 | NM_001081212.1 | 1479-2249 |
| Nfix | NM_001081981.3 | 477-1825 |
| Nr2f1 | NM_010151.2 | 642-1408 |
| Nr3c1 | NM_008173.3 | 416-1385 |
| Nrip1 | NM_173440.2 | 3395-3975 |
| Phlda1 | NM_009344.3 | 731-1366 |
| Rcn1 | NM_009037.2 | 711-1373 |
| Tle3 | NM_001083927.1 | 1980-2982 |
| Unc5b | NM_029770.2 | 2318-3161 |

**Data S1. Differential peaks for brain ERα CUT&RUN.** DiffBind output for ERα CUT&RUN experiment from ERα-expressing brain regions, along with ChIPSeeker gene annotations. Includes lists of genes annotated to brain-specific ERα and tissue-shared peaks.

**Data S2. Differential gene list for BNSTp ERα+ neuron RNA-seq.** DESeq2 comparisons and HT counts for RNA-seq experiments from BNSTp tissue.

**Data S3. Differential peaks for BNSTp ERα+ neuron ATAC-seq.** DiffBind output for ATAC-seq experiment from BNSTp, along with ChIPSeeker gene annotations.

**Data S4. Sexually dimorphic genes in each BNSTp snRNA-seq cluster.** DESeq2 output for differential expression analysis between male and female pseudo-bulk replicates within each BNST snRNA-seq cluster.

**Data S5. ERα target genes with male-biased expression in each BNSTp snRNA-seq cluster.** BETA output for ERα CUT&RUN peaks and male-biased genes in BNSTp clusters with significant over-representation of ERα binding.

**Data S6. Differential gene list for P4 BNST ERα+ nuclear RNA-seq.** DESeq2 output for differential expression analysis between neonatal female EB and neonatal female Veh groups. Additional list of neonatal EB-induced ERα target genes that are male-biased in adult BNSTp.

